# Three-dimensional synaptic organization of the human hippocampal CA1 field

**DOI:** 10.1101/2020.02.25.964080

**Authors:** Marta Montero-Crespo, Marta Domínguez-Álvaro, Patricia Rondón-Carrillo, Lidia Alonso-Nanclares, Javier DeFelipe, Lidia Blázquez-Llorca

## Abstract

The hippocampal CA1 field integrates a wide variety of subcortical and cortical inputs, but its synaptic organization in humans is still unknown due to the difficulties involved studying the human brain via electron microscope techniques. However, we have shown that the 3D reconstruction method using Focused Ion Beam/Scanning Electron Microscopy (FIB/SEM) can be applied to study in detail the synaptic organization of the human brain obtained from autopsies, yielding excellent results. Using this technology, 24,752 synapses were fully reconstructed in CA1, revealing that most of them were excitatory, targeting dendritic spines and displaying a macular shape, regardless of the layer examined. However, remarkable differences were observed between layers. These data constitute the first extensive description of the synaptic organization of the neuropil of the human CA1 region.

## Introduction

The hippocampus plays a crucial role in spatial orientation, learning and memory, and many pathological conditions (e.g., epilepsy and Alzheimer’s disease) are closely associated with synaptic alterations in the hippocampus (*1*). As has been previously discussed, one of the first steps towards understanding the way in which neuronal circuits contribute to the functional organization of the brain involves defining the brain’s detailed structural design and mapping its connection matrix (*2*). The connectivity of the brain can be examined at three major levels of resolution (*3*): (i) macroscopically, focusing on major tract connectivity; (ii) at an intermediate resolution, using light microscopy techniques that allow putative synaptic contacts to be mapped; and (iii) at the ultrastructural level, using electron microscopy (EM) to map true synaptic contacts. Numerous studies have described the ultrastructural characteristics and organization of hippocampal synapses in experimental animals (*4*). However, there is very little information about the synaptic organization of the human hippocampus and the brain in general, which is a major problem since the question remains as to how much of the animal model information can be reliably extrapolated to humans. The majority of these studies are performed in specimens removed during the course of neurosurgery in patients with tumors or intractable epilepsy (*5–10*). Since it is inevitable that surgical excisions pass through cortical regions that are normal, this represents an excellent opportunity to study human brain material. The problem is that although this material is thought to be close to what would be expected in the normal brain, the results cannot be unequivocally considered as representative of the normal condition of the human brain. Thus, a major goal in neuroscience is to directly study human brain with no recorded neurological or psychiatric alterations. In the present study, we started to address the issue of the hippocampal synaptic organization by focusing on the CA1 field. This hippocampal field receives and integrates a massive amount of information in a laminar-specific manner, and sends projections mainly to the subiculum and to extrahippocampal subcortical nuclei and polymodal association cortices (Fig. S1).

Studying the human brain via EM techniques presents certain problems and the scarcity of human brain tissue that is suitable for the study of synaptic circuitry is one of the most important issues to overcome. Recently, we have shown that the 3D reconstruction method using Focused Ion Beam/Scanning Electron Microscopy (FIB/SEM) can be applied to study in detail the synaptic organization of the human brain obtained from autopsies, yielding excellent results (*11, 12*).

For these reasons, we used FIB/SEM technology to perform a 3D analysis of the synaptic organization in the neuropil in all layers of the CA1 region from five human brain autopsies with a short postmortem delay. Specifically, we studied a variety of synaptic structural parameters including the synaptic density and spatial distribution, type of synapses, postsynaptic targets and the shape and size of the synaptic junctions.

The data reported in the present work constitutes the first extensive description of the synaptic organization in the human hippocampal CA1 field, which is a necessary step for better understanding its functional organization in health and disease.

## Results

We used coronal sections of the human hippocampus at the level of the hippocampal body and examined the CA1 field at both light and EM levels. Following a deep to superficial axis, the following main CA1 layers were analyzed: the alveus, *stratum oriens* (SO), *stratum pyramidale* (SP), *stratum radiatum* (SR) and *stratum lacunosum-moleculare* (SLM) (Fig. S2). Additionally, SP was subdivided into a deep part (dSP) close to the SO, and a superficial part (sSP), close to the SR.

### Light microscopy: volume fraction occupied by cortical elements

First, we estimated the total thickness of the CA1 field ―including the alveus ― in the radial axis. The average thickness was 2.70±0.62 mm. Following a deep-superficial axis, the average length of each layer was: 0.34±0.12 in the alveus; 0.06±0.03 mm in SO; 1.13±0.33 mm in SP; 0.55±0.31 mm in SR; and 0.62±0.16 mm in SLM. Thus, in relative terms, SP contributed the most to the total CA1 thickness (42%) followed by SLM (23%) then SR (20%), the alveus (13%) and SO (2%) (Fig. 1, Table S1). We then assessed the cellular composition of every CA1 layer, including the volume fraction (V_v_) occupied by different cortical elements (i.e., blood vessels, glial and neuronal somata and neuropil), estimated by applying the Cavalieri principle (*13*). The neuropil constituted undoubtedly the main element in all layers (more than 90%; Fig. S3b,f, Table S1) followed by blood vessels (range from 4.79% in SR to 7.58% in SO; Fig. S3b,c, Table S1). The volume fraction occupied by glial cell and neuronal bodies was less than 2% (Fig. S3b,d,e, Table S1), except for SP, where neuronal cell bodies occupied a volume of 4.23±1.07% (Fig. S3, Table S1). As expected, the volume occupied by neurons was significantly higher in SP than in any other layer (ANOVA, p<0.001). The neuropil was significantly more abundant in SR (94.19±1.17%) than in SP (90.11±1.32%, ANOVA, p=0.015) and SO (90.01±3.07%; ANOVA, p=0.012). No further significant differences regarding cortical elements were found between any other layers.

**Fig. 1.**
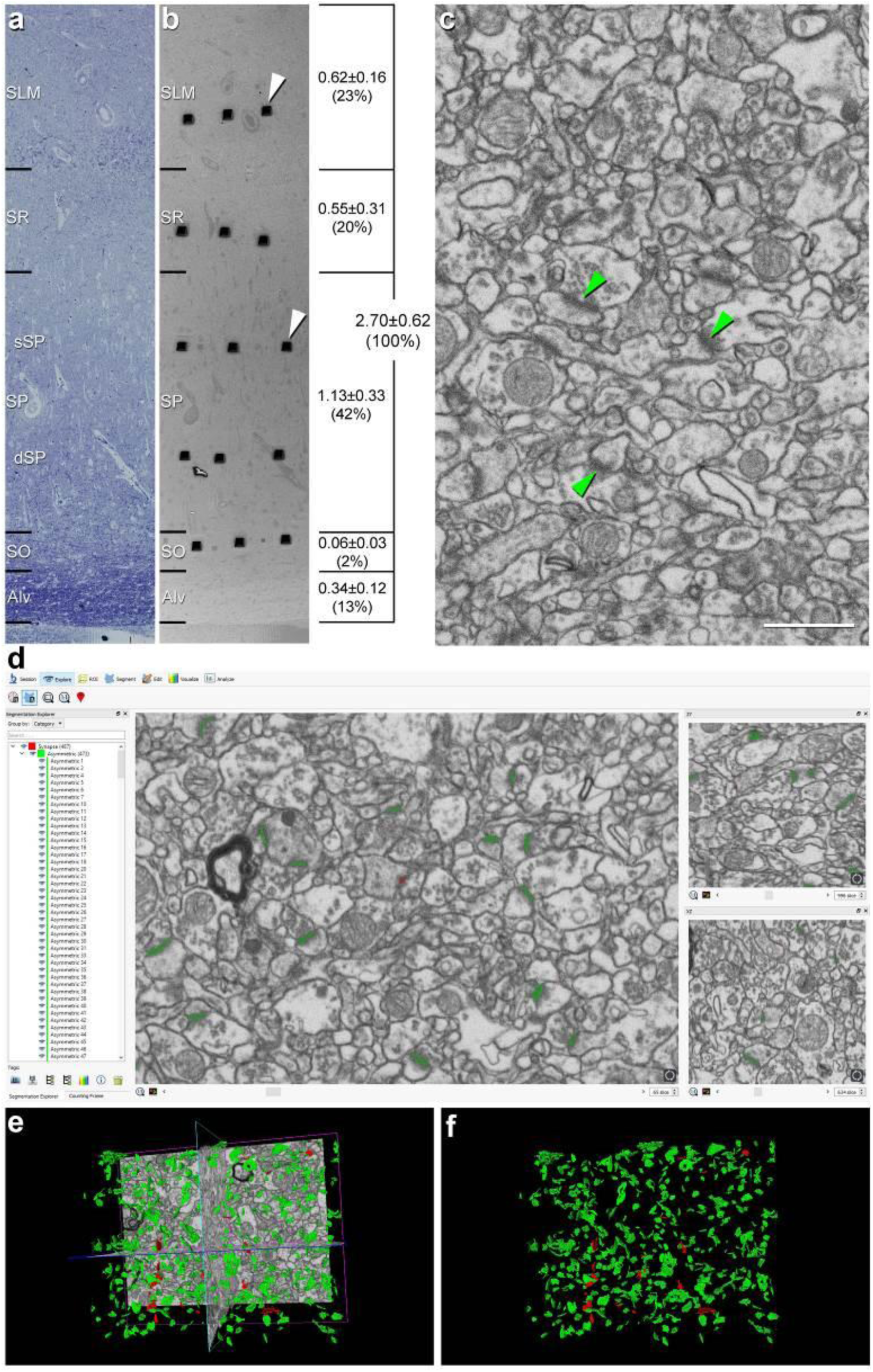
Correlative light/electron microscopy analysis of CA1 using FIB/SEM and EspINA software. **a, b,** Delimitation of layers is based on the staining pattern of 1 µm thick semithin section stained with toluidine blue (**a**). This section is adjacent to the block surface (**b**), which is visualized with the SEM. This allows the exact location of the region of interest to be determined. The thickness of each stratum (mm; mean±SD), as well as its relative contribution to the total CA1 thickness, is shown on the right side of panel **b**. White arrowheads in **b** point to two of the trenches made in the neuropil (three per layer). **c**, FIB/SEM image at a magnification of 5 nm/pixel. Some asymmetric synapses have been marked with green arrowheads. **d**, Screenshot from the EspINA software interface. The stacks of images are visualized with EspINA software, permitting the identification and 3D reconstruction of all synapses in all spatial plans (XY, XZ and YZ). **e**, Shows the three orthogonal planes and the 3D reconstruction of segmented synapses. **f**, Only the segmented synapses are shown. AS are colored in green and SS in red. Scale bar in **c** corresponds to: 170 µm in a–b; 1 µm in **c**

### Electron microscopy

#### Distribution of synapses in the neuropil

Each single reconstructed synapse was sorted according to different qualitative and quantitative parameters (see Material and Methods). Specifically, regarding qualitative characteristics, we distinguished four different parameters: i) the type of synapses: asymmetric synapses (AS) or symmetric synapses (SS); ii) the postsynaptic targets: axospinous (on the head or neck of the dendritic spine) or axodendritic (on spiny or aspiny dendritic shafts); and iii) the synaptic shape: macular, horseshoe-shaped, perforated or fragmented synapses. Additionally, three quantitative parameters were used for classification: i) the synaptic apposition surface (SAS) area, ii) SAS perimeter and iii) SAS curvature.

##### Synaptic density

All synapses (n=24,752) in the 75 stacks of images examined were fully reconstructed. After discarding the synapses not included in the unbiased counting frame (CF), a total of 19,269 synapses (AS=18,138; SS=1,131) were further considered for analysis and classification. The number of synapses per volume unit in every layer was calculated (synaptic density). The mean synaptic density was 0.67±0.21 synapses/µm^3^ (Table 1). Differences in synaptic density between layers were observed (Fig. 2a; Table 1); sSP was the layer with the highest number of synapses per volume unit (0.99±0.18 synapses/µm^3^), whereas SO had the lowest synaptic density (0.45±0.19 synapses/µm^3^). However, synaptic density differences were only statistically significant between sSP and both SO (ANOVA, p=0.0005) and SLM (0.52±0.08 synapses/µm^3^; ANOVA, p=0.002; Fig. 2a).

**Fig. 2.**
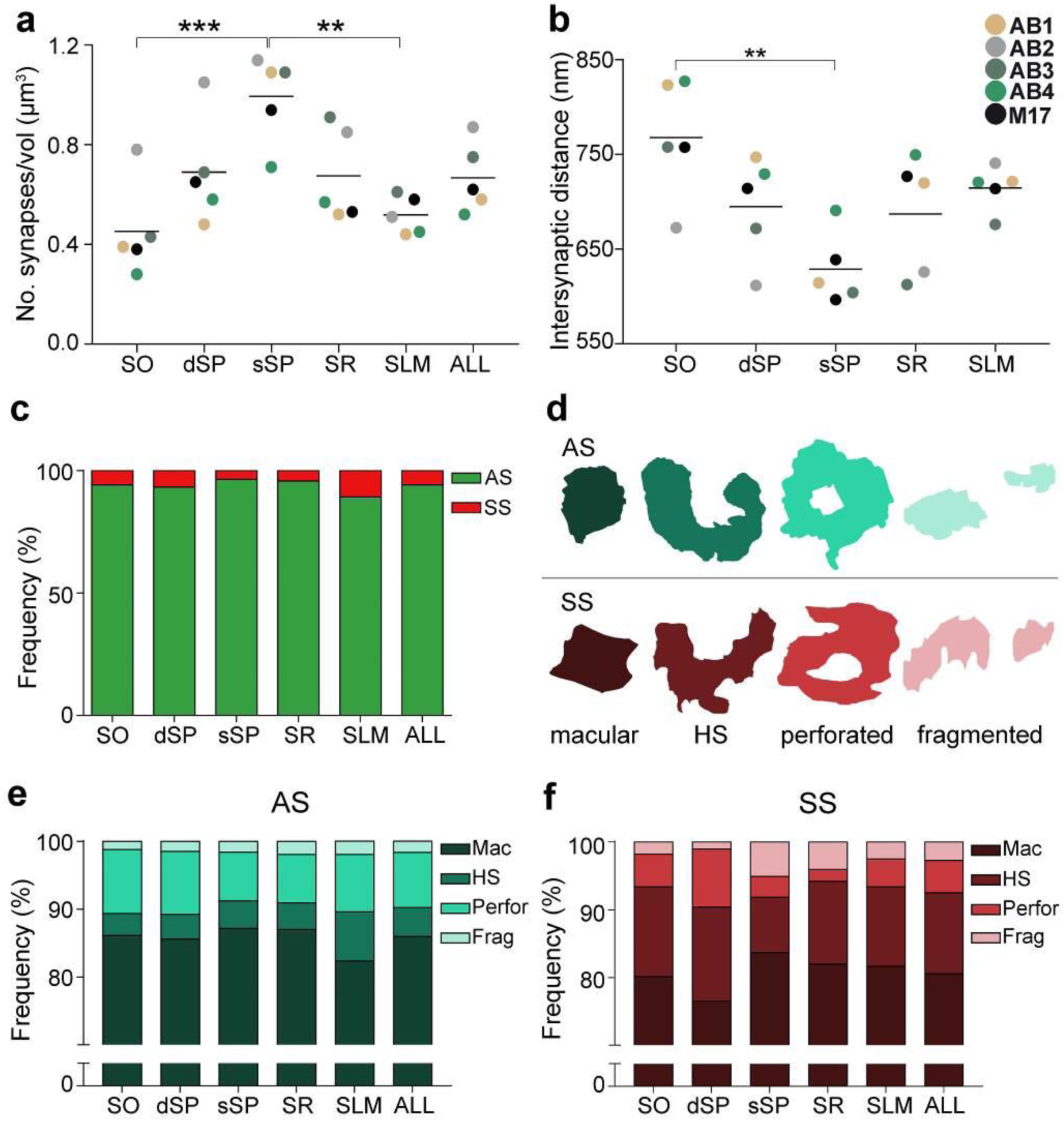
Synaptic density, intersynaptic distance, proportion of asymmetric synapses (AS) and symmetric synapses (SS), and proportion of synaptic shapes in CA1. **a**, Graph showing the mean synaptic density in all layers. **b**, Graph showing the mean intersynaptic distance in all layers. Each dot in **a** and **b** represents the data from each case, with the grey line showing the mean value. **c**, Shows the percentages of AS and SS in all layers. **d**, Illustrates examples of the different types of synapses based on the shape of the synaptic junction: macular, horseshoe-shaped (HS), perforated and fragmented. The upper and lower rows show examples of shapes of AS and SS, respectively. **e, f**, Percentages of the different types of synaptic shapes within the population of AS (**e**) and SS (**f**) in all layers.

**Table 1.** Data regarding synapses in all layers of the CA1. CF: counting frame; SD: standard deviation.

|  | SO | dSP | sSP | SR | SLM | All layers |
| --- | --- | --- | --- | --- | --- | --- |
| No. AS | 2648 | 3849 | 5183 | 3836 | 2622 | 18138 |
| No. SS | 166 | 281 | 196 | 172 | 316 | 1131 |
| No. synapses<br>(AS+SS) | 2814 | 4130 | 5379 | 4008 | 2938 | 19269 |
| % AS | 94.10% | 93.20% | 96.36% | 95.71% | 89.24% | 94.13% |
| % SS | 5.90% | 6.80% | 3.64% | 4.29% | 10.76% | 5.87% |
| CF volume ($\mu\text{m}^3$) | 6221 | 6004 | 5400 | 6007 | 5690 | 29322 |
| No. AS/$\mu\text{m}^3$<br>(mean±SD) | 0.43±<br>0.19 | 0.64±<br>0.22 | 0.96±<br>0.18 | 0.64±<br>0.19 | 0.46±<br>0.07 | 0.63±<br>0.21 |
| No. SS/$\mu\text{m}^3$<br>(mean±SD) | 0.03±<br>0.01 | 0.05±<br>0.02 | 0.04±<br>0.01 | 0.03±<br>0.01 | 0.06±<br>0.02 | 0.04±<br>0.01 |
| No. all synapses/$\mu\text{m}^3$<br>(mean±SD) | 0.45±<br>0.19 | 0.69±<br>0.22 | 0.99±<br>0.18 | 0.67±<br>0.19 | 0.52±<br>0.08 | 0.67±<br>0.21 |
| Intersynaptic distance<br>(nm; mean±SD) | 742.81±<br>63.06 | 669.81±<br>54.18 | 604.00±<br>38.08 | 653.77±<br>69.51 | 689.65±<br>23.72 | - |

##### Spatial distribution

Synapses fitted into a random spatial distribution in all layers since the observed F, G and K functions laid within the envelope generated by 99 simulations of the CSR model (*14, 15*) (Fig. S4).

Furthermore, significant differences in the average intersynaptic distance were only found between sSP (604.00±38.08 nm) and SO (742.81±63.06 nm, ANOVA, p=0.0027; Fig. 2b; Table 1). The maximum value was found in SO, whereas the minimum value was observed in sSP (Fig. 2b; Table 1). Moreover, the variables synaptic density and intersynaptic distance were strongly and indirectly correlated (R^2^=0.90).

##### Proportion of AS and SS

It is well established that AS are mostly glutamatergic and excitatory, whereas SS mostly GABAergic and inhibitory (*16*) (Fig. 3). Therefore, the proportions of AS and SS were calculated in each layer. Since synaptic junctions were fully reconstructed in the present study, all of them could be classified as AS or SS based on the thickness of their PSDs (*17*) (Fig. 3).

**Fig. 3.**
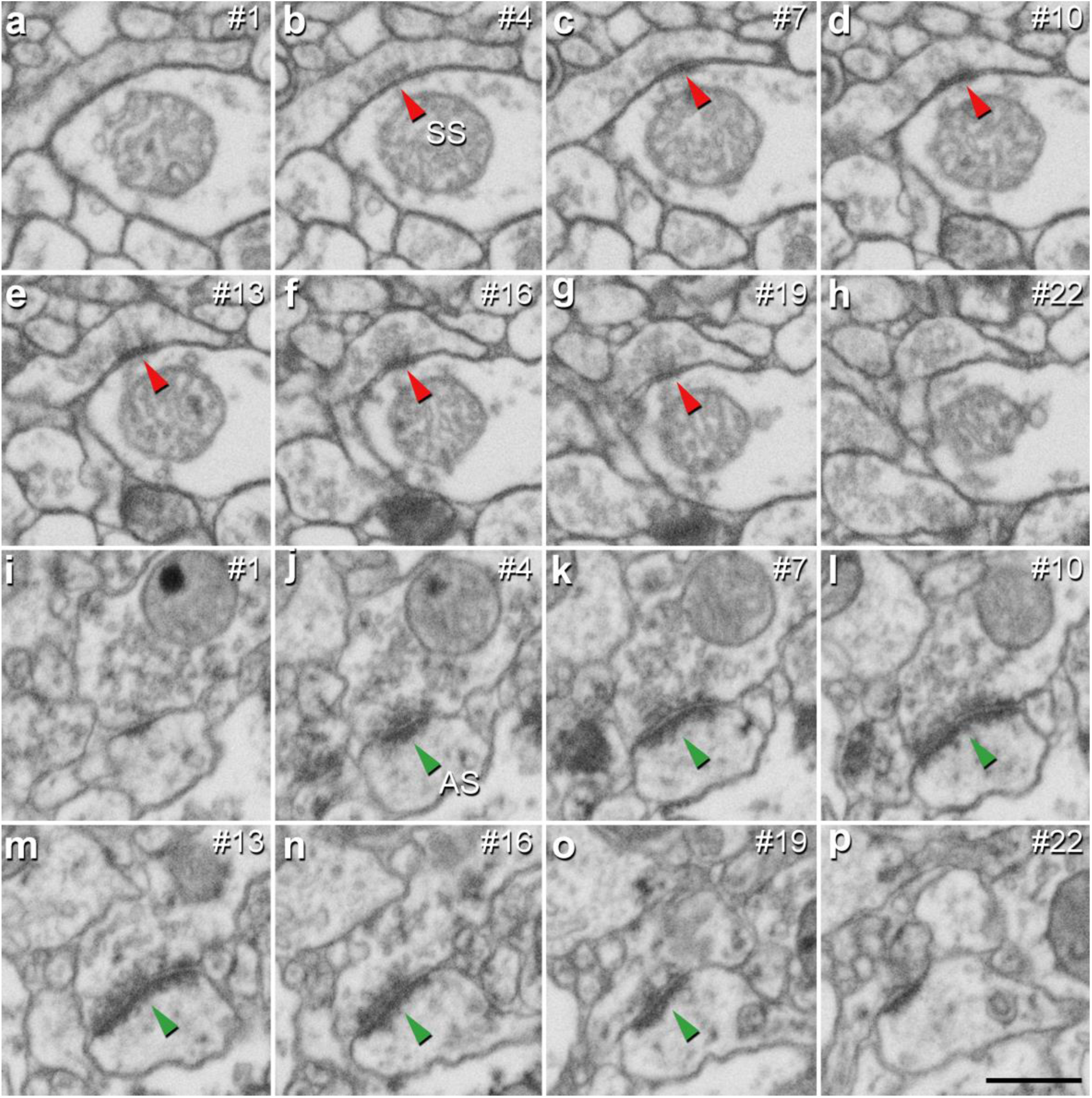
Identification of an asymmetric synapse (AS) and a symmetric synapse (SS) in the neuropil of the human CA1 region. **a–h,** Crops from an electron microscopy section obtained by FIB/SEM to illustrate an SS (red arrowhead). **i–p,** Crops from electron microscopy images following an AS (green arrowhead). The number of the section is indicated in the top right hand corner of each section, with a 60 nm thickness separation between images. Synapse classification was based on the examination of the full sequence of serial material. Scale bar in **p** corresponds to: 500 nm in **a–p**.

The AS:SS ratio was close to 95:5 in all layers, except SLM, where the percentages were close to 90:10 (Fig. 2c; Table 1). We found significant differences in the proportion of excitatory and inhibitory contacts between layers (χ^2^, p<0.0001). Specifically, the frequency of AS was significantly lower in dSP (93.20%) as compared to sSP (96.36%; χ^2^, p=4.381×10^-12^) and SR (95.71%; χ^2^, p=7.478×10^-7^; Fig. 2c; Table 1). Furthermore, the proportion of SS was significantly higher in SLM than in any other layer (10.76%; χ^2^, p<0.0001, Table 1).

##### Postsynaptic targets

Two main postsynaptic targets were considered (Fig. 4): dendritic spines (axospinous synapses) and dendritic shafts (axodendritic synapses). In the case of axospinous synapses, the exact location of the synaptic contact was determined (i.e., the head or neck of the dendritic spine, Fig. 4a–k). For axodendritic synapses, dendritic shafts were further classified as spiny (when dendritic spines could be observed emerging from the shaft) or aspiny. Only synapses whose postsynaptic target was clearly identifiable after navigation through the stack of images (n=9,442; AS = 8,449, SS = 993) were considered for analysis.

**Fig. 4.**
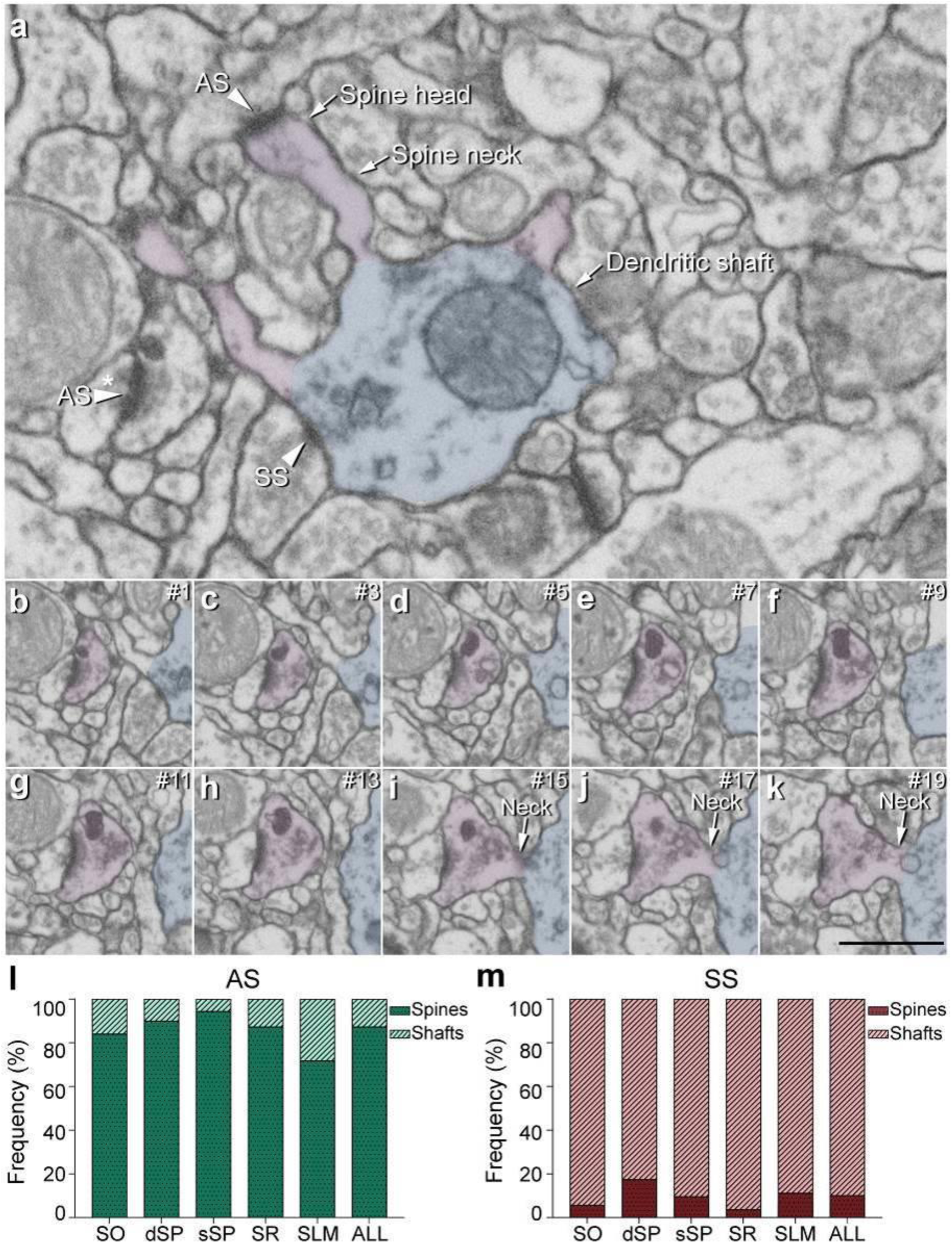
Postsynaptic target identification in serial electron microscopy images. **a**, A crop from an electron microscopy section to illustrate a dendritic shaft (blue) with three dendritic spines (purple) emerging from the shaft (the neck and head have been indicated in one of the spines). An SS on the dendritic shaft is pointed out with an arrowhead. An axospinous AS (marked with an arrowhead) is established on the head of one of the spines. Another AS is indicated (arrowhead with asterisk); however, the nature of the postsynaptic element where the synapse is established cannot be distinguished in a single section. **b–k**, Crops from electron microscopy serial sections to illustrate the nature of the postsynaptic element of the AS (arrowhead with asterisk) in **a**. By following up from this AS through the stack of images (the number of the section is indicated in the top right hand corner of each section; 40 nm thickness separation between images), a dendritic spine (purple), whose neck has been labeled emerging from the dendritic shaft (blue), can be unequivocally identified. **l, m**, The percentage of axospinous and axodendritic synapses within the AS (**l**) and SS (**m**) populations in all layers of CA1. Scale bar in **k** corresponds to: 1 µm in **a**; 500 nm in **b–k**.

#### Total synaptic population

Despite the great disparity between layers, most synapses (AS+SS) were established on dendritic spines —especially on the head— (n=7,469; 79.10%, ranging from 59.12% in SLM to 88.29% in sSP, Table S2), rather than on dendritic shafts (n=1,973; 20.90%, ranging from 11.71% in sSP to 40.88% in SLM, Table S2). Synapses (AS+SS) on spiny shafts were more abundant than synapses on aspiny shafts in all CA1 layers, except for SLM (χ^2^, p<0.0001, Table S2).

As a whole, axospinous AS were clearly the most abundant type of synapses in all layers (n=7,369; 78.04%, ranging from 56.80% in SLM to 87.61% in sSP, Figs. 5, S5; Table S2), followed by axodendritic AS, except for sSP (n=1,080; 11.44%, ranging from 5.25% in sSP to 22.42% in SLM; Figs. 5,S5; Table S2), where axodendritic SS were the second most abundant type of synapses (n=893; 9.46%, ranging from 6.46% in sSP to 18.46% in SLM; Figs. 5,S5; Table S2). Axospinous SS were remarkably scarce (n=100; 1.06%, ranging from 0.37% in SR to 2.32% in SLM; Figs. 5, S5; Table S2).

**Fig. 5.**
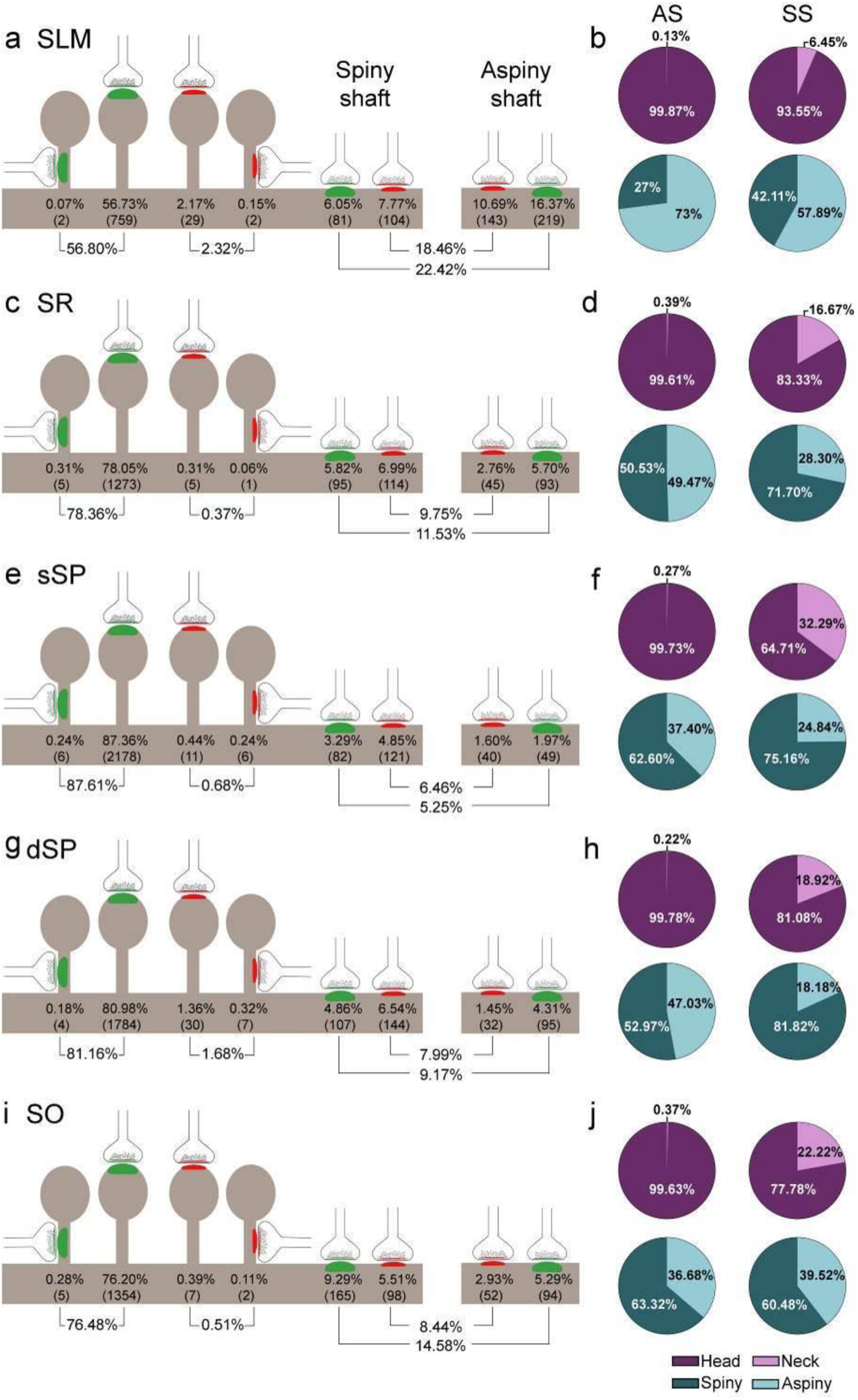
Representation of the distribution of synapses according to their postsynaptic targets in all layers of CA1. **a, c, e, g, i**, Shows the percentages of axospinous (both on the head and the neck of dendritic spines) and axodendritic (both on spiny and aspiny shafts) asymmetric synapses (AS;green) and symmetric synapses (SS;red). The numbers of each synaptic type are shown in brackets. **b, d, f, h, j**, Pie charts to illustrate the proportions of AS and SS according to their location as axospinous synapses (i.e., on the head or on the neck of the spine) or axodendritic synapses (i.e., spiny or aspiny shafts).

Significant differences in the proportion of synapses were found between layers (χ^2^, p<0.0001) (Table S3). Both axodendritic AS and axodendritic SS were clearly more frequent in SLM than in any other layer (χ^2^, p<0.0001). sSP presented the largest proportion of axospinous AS (χ^2^, p<0.0001) and the lowest frequency of axodendritic AS (χ^2^, p<0.0001). Additionally, a lower prevalence of axospinous AS and a larger proportion of axodendritic AS were observed in SO compared to dSP (χ^2^, p=0.0004 and p=1.518×10^-7^, respectively) (Tables S4–S7). Finally, the prevalence of axospinous SS was significantly higher in dSP and SLM than in any other layer (χ^2^, p<0.001 in dSP vs SO and sSP; p<0.0001 in the rest of the cases).

#### Postsynaptic preference of AS and SS

Regardless of the layer, most AS were established on dendritic spines (n=7,369; 87.22%, ranging from 71.70% in SLM to 94.34% in sSP in the population of AS; Fig. 4l; Table S8), and they were found almost exclusively on the head of the spines (>99.5% in all layers, Fig. 5). The remaining AS were established on dendritic shafts (n=1,080; 12.78%, ranging from 5.66% in sSP to 28.30% in SLM; Fig. 4l; Table S8), with a preference for spiny shafts in SO and sSP, whereas in SLM the preference was for aspiny shafts (Fig. 5). In the case of SS, most were axodendritic (n=893; 89.93%), ranging from 82.63% in dSP to 96.36% in SR (Fig. 4m; Table S8). SS showed a clear preference for spiny shafts in all layers except in SLM (Fig. 5). The remaining SS were established on dendritic spines (n=100; 10.07%, ranging from 3.64% in SR to 17.37% in dSP; Fig. 5m; Table S8). These axospinous SS were found especially on the head of the spines (82%, Fig. 5).

In every layer, we found a consistent association for AS and dendritic spines, and for SS and dendritic shafts (χ^2^, p<0.0001). Moreover, the preference of inhibitory contacts for dendritic shafts was found for both spiny and aspiny dendritic shafts, regardless of the layer (χ^2^, p<0.0001). Spiny shafts received a higher proportion of SS than AS in all layers, except for SO, while aspiny shafts received a higher proportion of AS than SS in all layers, especially in dSP. Although scarce, dendritic spines receiving multiple synapses were found in all layers (2.11% of total spines in all layers; Fig. S5; Table S9), whereas single axospinous SS were extremely rare or even not found in some layers (Fig S5; Table S9). Moreover, multiple-headed dendritic spines (double-headed in most cases) were also observed (1.39% of total spines in all layers; Fig S5; Table S9).

##### Shape of the synaptic junctions

Synapses were categorized as macular, horseshoe-shaped, perforated or fragmented (n=19,269; AS=18,138, SS=1,131; Fig. 2d–f). The vast majority of both AS and SS (more than 75% in all layers) had a macular shape (85.95% and 80.55%, respectively; Fig. 2e–f; Table S10), followed by perforated synapses in the case of AS (8.13%, Fig. 2e–f; Table S10) and horseshoe-shaped synapses in the case of SS (11.94%, Fig. 2e–f; Table S10). We observed that some synaptic shapes were more prevalent in some layers. Overall, AS with complex shapes (that is, including either horseshoe-shaped, perforated or fragmented) were more abundant in SLM than in any other layer (χ^2^, p<0.001; Fig. 2e), especially horseshoe-shaped synapses (χ^2^, p<0.0001; Fig. 2e). They were mainly located in dendritic shafts (χ^2^, p=1.259×10^-5^; Fig. S6; Table S11). Additionally, perforated AS were observed more frequently in SO and dSP than in sSP and SR (χ^2^, p<0.001; Fig. 2e). No differences could be observed in the case of SS (χ^2^, p>0.001).

Considering both AS and SS against the four types of synaptic shapes in each layer, we found that horseshoe-shaped synapses were significantly more abundant in the SS population than in the AS population in all layers (χ^2^, p<0.001). No synaptic shape was more frequent among AS.

#### Size of the synapses

Morphological features of SAS were extracted with EspINA software for both AS and SS (n=19,269; AS=18,138, SS=1,131; Figs. 6,S7–S8; Tables S12–S14).

**Fig. 6.**
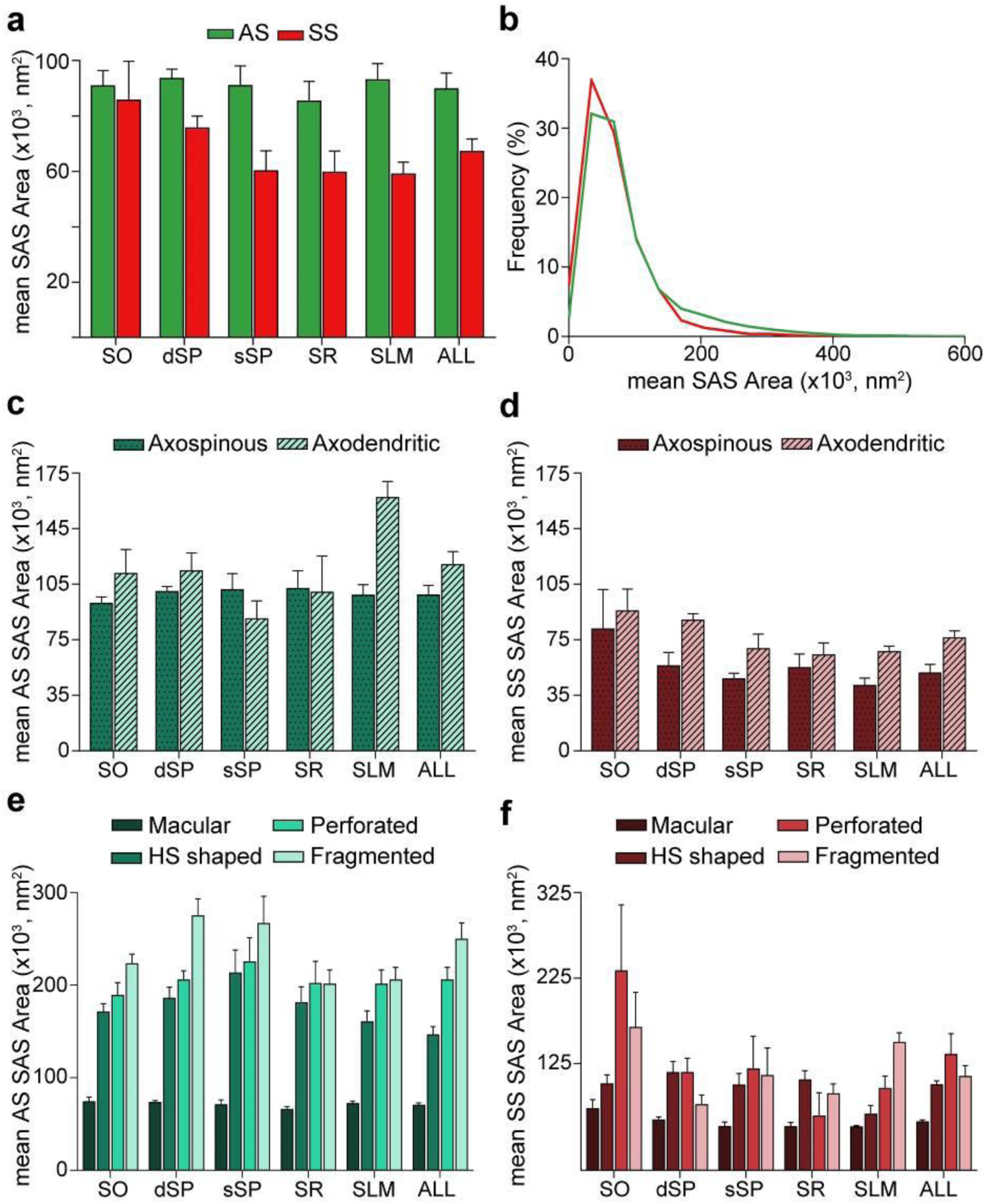
Synaptic surface area (SAS) measurements. **a,** Mean SAS area of asymmetric synapses (AS;green) and symmetric synapses (SS;red) are represented for each layer of CA1 (mean±sem). **b**, Frequency distribution of SAS areas for both AS (green line) and SS (red line) in all layers of CA1. No differences were observed in the frequency distribution of SAS areas between the two synaptic types (KS, p>0.05). **c**, **d**, Mean SAS area of axospinous and axodendritic synapses are also shown for AS (**c**) and SS (**d**) in the whole CA1 and per layer (mean±sem). Both axodendritic AS and SS were larger in SLM than axospinous AS (MW, p<0.01) and SS (MW, p<0.05), respectively, while in the rest of the layers, no differences were observed. **e**, **f**, Mean SAS area related to the different synaptic shapes are plotted for both AS (**e**) and SS (**f**) in all layers of CA1 (mean±sem). Macular synapses are significantly smaller than the other more complex-shaped ones (i.e., HS, perforated and fragmented); however, this difference is only significant for AS (ANOVA, p<0.0001).

The mean SAS areas of AS and SS were 89,727.65 nm^2^ and 67,236.17 nm^2^, respectively, while the mean SAS perimeters of AS and SS were 1,458.82 and 1,378.38 nm, respectively (Table S12). No differences were observed between layers regarding the size (area and perimeter) of the synapses both for AS (ANOVA, p>0.05; Table S12) and SS (ANOVA, p>0.05; Table S12). However, AS had significantly larger areas than SS when considering all synapses together (MW, p=0.032; Fig. 6a; Table S12), but when focusing on particular layers, this difference in area between AS and SS was only observed in dSP, sSP and SLM (MW, p=0.016, p=0.032 and p=0.008, respectively; Fig. 6a; Table S12). No differences were found in perimeter measurements (MW, p>0.05). Although significant differences in SAS area did not extrapolate to differences in SAS perimeter, there was a strong correlation between these two parameters (R^2^ = 0.81 for all synapses; R^2^ = 0.82 for AS; R^2^ = 0.81 for SS).

To further characterize the size distribution of the SAS of both AS and SS, we plotted the frequency histograms of SAS areas for each individual layer and all layers. Frequency histograms had similar shapes for both types of synapses when considering all layers and within each layer, with a positive skewness (that is, most synapses presented small SAS area values). Moreover, the frequency distributions of AS and SS greatly overlapped, as did the frequency distributions of SAS area between the layers (KS, p>0.001; Fig. 6b, Fig. S7).

Additionally, we studied synaptic size regarding the postsynaptic targets. Although both the mean SAS area and the perimeter of axodendritic AS were larger (117,360.02 nm^2^ and 1,686.99 nm, respectively) than axospinous AS (98,200.61 nm^2^ and 1,548.38 nm, respectively), these differences were not statistically significant (MW, p>0.05; Fig. 6c; Table S13). Only axodendritic AS in SLM were significantly larger than axospinous AS regarding both SAS area and perimeter (MW, p=0.008; Fig. 6c; Table S13). Overall, axodendritic SS (71,218.23 nm^2^) had a significantly larger mean area than axospinous SS (49,044.59 nm^2^) (MW, p=0.032; Fig. 6c; Table S13) but, again, this difference in the mean SAS area was significant only in SLM (MW, p=0.032; Fig. 6d; Table S13).

Finally, analyses were carried out to determine the differences in synaptic size in terms of the shape of the synaptic junctions. Macular synapses were smaller than the rest of the more complex-shaped synapses for both AS (mean macular SAS area: 70,322.92 nm^2^, mean complex-shaped SAS area: 200,539.32 nm^2^) and SS (mean macular SAS area: 56,769.81 nm^2^, mean complex-shaped SAS area: 115,170.07 nm^2^). However, these differences were only significant in the case of AS, as demonstrated by both mean SAS area and perimeter (ANOVA, p<0.0001 in all cases, except for macular AS and horseshoe-shaped AS, p=0.002; Fig. 6e-f; Table S14). This difference was also observed between AS in all layers (ANOVA, p<0.05; Fig. 6e; Table S14). No differences were observed in the synaptic size of the different synaptic shapes between the layers (ANOVA, p>0.05).

See Supplementary materials for information about SAS Curvature.

### Interindividual variability

Differences between cases were observed regarding several of the parameters examined in several layers (Tables S15-S30). All significant differences are reported under the corresponding tables for each individual case. Importantly, differences between individual cases were not necessarily found with respect to the same parameter, or in the same layer or in the same direction (increase or decrease). For instance, case AB1 presented a larger volume fraction of blood vessels in SO than the rest of the cases (ANOVA, p<0.05; Table S15), except for case AB2. In case AB2, the volume fraction occupied by neuronal bodies in SR was higher than in the rest of the subjects, except for AB3 (ANOVA, p<0.01; Table S15). Additionally, the volume occupied by glia in the SLM of case AB1 was higher than in the rest of the cases, except for AB3 (ANOVA, p<0.05; Table S15). Furthermore, compared to the rest of the cases, AB2 and AB3 presented higher synaptic densities in SR (ANOVA, p<0.05; Table S19). Also, compared to the rest of the subjects, AB2 exhibited a higher synaptic density (ANOVA, p<0.01; Table S16). In addition, the proportion of SS was higher in SLM in M17 than in AB1, AB2 and AB3 (χ^2^, p<0.001; Table S20).

When focusing on postsynaptic targets, SLM was one of the layers with the greatest differences among cases (χ^2^, p=1.000×10^-17^; Table S25). In this layer, out of all the cases, case AB1 exhibited the highest proportion of axospinous AS and the lowest percentage of both axodendritic AS and SS (χ^2^, p<0.0001; Table S25). Additionally, a larger proportion of axospinous AS was also observed in subject AB3 when compared to cases AB2 and M17 (χ^2^, p<0.0001; Table S25).

Macular synaptic junctions were clearly the most abundant type in all cases and layers. However, perforated AS were especially abundant in subject AB4 compared to the rest of the cases in all layers (χ^2^, p<0.001; Table S26-S30) with the exception of AB1 in sSP. Additionally, in case AB4, the AS in sSP were larger than in the rest of the individuals (ANOVA, p<0.0001; Table S18), apart from in the case of M17.

## Discussion

The present study constitutes the first exhaustive description of the synaptic organization in the neuropil of the human CA1 field using 3D EM. The following major results were obtained: (i) there are significant differences in the synaptic density between layers; (ii) synapses fitted into a random spatial distribution; (iii) most synapses are excitatory, targeting dendritic spines and displaying a macular shape, regardless of the layer — although significant differences were observed between certain layers; (iv) SLM showed several peculiarities compared with other layers, such as a larger proportion of inhibitory synapses, a higher prevalence of both AS and SS axodendritic synapses, and the presence of more complex synaptic shapes. The wide range of differences in the synaptic organization of the human CA1 layers found in the present study may be related to the variety of inputs arriving in a layer-dependent manner (Fig. S1).

### CA1 structural composition

The neuropil represents the main structural component of CA1 (more than 90% of all layers). The contribution of SP to the total CA1 radial extension accounted for almost a half of the total thickness (SP thickness: 1.13 mm; total CA1 thickness: 2.70 mm). This great extent of SP represents a major difference with the rodent brain and other species. Indeed, important variances can be observed in the hippocampal neuroanatomy of humans compared to rodents (*1,18-21*). In the rat hippocampus, SP is around five cell bodies thick and neuronal somata are densely packed, being SR the layer that contributes the most to the total CA1 thickness. In humans, SP can be up to 30 cell somata thick, with a wider separation of neurons compared to other species. As previously discussed in (*18*), this sometimes refers to a “corticalization” of the human CA1 pyramidal cell layer because it resembles a neocortical cytoarchitecture, which most probably has fundamental functional and hodological consequences: the basal and apical dendrites of human pyramidal cells are intermixed in the pyramidal cell layer (Fig. S1), whereas in rodents, the basal and apical dendritic arbors are basically separated (basal dendrites in SO; apical dendrites in SR).

### Synaptic density

Synapses were found in all layers except the alveus where they were virtually nonexistent. The mean synaptic density was 0.67 synapses/µm^3^. However, synaptic density was not homogenous among layers. Consistently in all individuals, the highest value was found in sSP (0.99 synapses/µm^3^), followed by dSP (0.69 synapses/µm^3^), while the lowest was observed in SO (0.45 synapses/µm^3^). Since no quantitative 3D analysis of the synaptic organization in the human hippocampus has been performed before, our data could not be compared to previous reports. However, in recent studies from our group using FIB/SEM to analyze the synaptic density in the rodent CA1 field, the following values were obtained: 2.53 synapses/µm^3^ in SO, 2.36 synapses/µm^3^ in SR and 1.72 synapses/µm^3^ in SLM in the mouse (*22*), and 2.52 synapses/µm^3^ in SR in the rat (Blazquez-Llorca et al., in preparation). These values are much higher than the ones found in the present work for the human CA1 (Table 1). Such huge differences in synaptic density between humans and rodents —together with the above-mentioned divergences in the morphology and distribution of pyramidal cells in the SP of CA1 (*18*), as well as differences in other anatomical, genetic, molecular and physiological features (*18,21-25*)— further support the notion that there are remarkable differences between the human and rodent CA1. These differences clearly need to be taken into consideration when making interpretations in translational studies comparing one species to another.

### Spatial synaptic distribution, proportion of synapses and postsynaptic targets

While synaptic density differed across layers, the spatial organization of synapses was consistently random in all layers. Randomly distributed synapses have also been described in the somatosensory cortex of rats and the frontal and transentorhinal cortices of the human brain (*11,15,26,27*), suggesting that this synaptic characteristic is a widespread ‘rule’ of the cerebral cortex of different species.

It has also been consistently reported that the neuropil is characterized by a much higher number of excitatory contacts compared to inhibitory synapses in different brain regions and species (4,*11,27–30*). In the present study, the density of inhibitory synapses was particularly low in most CA1 layers (AS:SS ratio in all layers was around 95:5 except for in SLM, where the ratio was close to 90:10). This data is in line with our study using FIB/SEM to analyze the synaptic density in the mouse (where the proportion of synapses that were inhibitory was 8% in the SLM, and approximately 2% in the case of the SR and SO) (*22*) and in the rat CA1 field (where 4% of the synapses in SR were inhibitory; Blazquez-Llorca et al., in preparation).

Regarding postsynaptic preferences, we observed a clear preference of excitatory axons and inhibitory axons for dendritic spines and dendritic shafts, respectively, which is also characteristic in other cortical regions and species, although variations in their percentages have been reported (*4,12,27,28,30-32*). For example, axospinous AS are especially abundant in sSP (87.61%) when compared to other brain regions in both humans and other species such as layer II of the human transentorhinal cortex, where axospinous AS account for only 55% of the total synaptic population (*12*).

### Shape and size of the synapses

Most synapses presented a simple, macular shape (accounting for 86% of the synapses in all layers of CA1), in agreement with previous reports in different brain areas and species (*12*,*33-36*).

The shape and size of the synaptic junctions are strongly correlated with release probability, synaptic strength, efficacy and plasticity (*37–40*). In this regard, all three types of non-macular synapses (with more complex shapes) were larger than macular ones. Although the functional significance of perforations is still unclear, perforated synapses are known to have more AMPA and NMDA receptors than macular synapses and are thought to constitute a relatively powerful population of synapses with more long-lasting memory-related functionality than their smaller, macular counterparts (*38,39,41*).

Considering all synapses, excitatory contacts were larger than inhibitory ones, as has also been observed in layer II of the human transentorhinal cortex (*11*); however, this contrasted with the findings in the somatosensory cortex (*36*) and SR of CA1 in the rat (Blazquez-Llorca et al., in preparation). A tendency towards axodendritic synapses being bigger than axospinous synapses was also observed; however, this difference was only significant in the case of SLM synapses. Complex-shaped AS were also found more frequently associated with axodendritic AS than with axospinous AS in SLM, while the opposite was the case for the rest of the layers. These findings agree with reports in the rat hippocampus, where excitatory synapses on SLM dendrites were observed to be: (i) larger than synapses in other layers; (ii) more frequently perforated (approximately 40%); and (iii) located to a greater extent on dendritic shafts (*30*).

### Relation between synaptic inputs and synaptic organization of each layer

The wide range of differences in the synaptic organization of the human CA1 layers found in the present study, especially between SLM and the rest of layers, may be related to the variety of inputs arriving to these layers (Figure S1). Unfortunately, detailed hippocampal human connectivity is far to be known: data directly obtained from human brains are very scarce and most data are inferred from rodents and primates (*42, 43*). In the primate brain, the CA1 field receives a wide variety of inputs from multiple subcortical and cortical brain regions (*42, 43*), being the major input to CA1 originated in the EC. Specifically, neurons located in layer III (and layer V) of the EC project directly to SLM, whereas neurons in layer II project to the rest of CA1 layers indirectly via the DG and CA3 field (*42, 44*). Considering both the synaptic data obtained in the present study and the connectivity knowledge in monkeys, it may seem that the synaptic organization in the layers receiving CA3 Schaffer collateral inputs (i.e. SO, SP and SR) differ with the synaptic organization found in the layer receiving direct inputs from the EC (i.e. SLM). Additionally, SLM receives a higher number of glutamatergic inputs from the amygdala and from the parietal and medial temporal cortex and higher numbers of 5-HT and Substance-P immunoreactive fibers, with a possible extrinsic origin in the Raphe nuclei and the laterodorsal tegmental nucleus (Figure S1).

It has been proposed that the CA3-CA1 synaptic connection plays a key role in the learning-induced synaptic potentiation of the hippocampus (*45*), while the direct projection from EC to SLM of CA1 seems to modulate information flow through the hippocampus (*46*). It has been reported that a high-frequency stimulation in SLM evokes an inhibition sufficiently strong to prevent CA1 pyramidal cells from spiking in response to Schaffer collaterals input (*46*). This finding could be supported by our present data showing an elevated inhibitory synapse ratio in comparison to other CA1 layers. It has also been described that afferences from the EC contact not only the apical tuft of CA1 pyramidal cells, but also interneurons of the SLM (*47*). One of these interneurons are the neurogliaform cells, which receive monosynaptic inputs from the EC and are also synaptically coupled with each other and with CA1 pyramidal cells (*48*). Whether the higher proportion of axodendritic synapses —particularly in aspiny shafts, which are likely to be originated from interneurons —, found in the present study in SLM compared to other CA1 layers is related to a particular synaptic circuit organization involving certain types of interneurons located in this layer remains to be elucidated.

## Online Methods

### Sampling procedure

Human brain tissue was obtained from autopsies (with short post-mortem delays of less than 4.5 hours) from 5 subjects with no recorded neurological or psychiatric alterations (supplied by Unidad Asociada Neuromax, Laboratorio de Neuroanatomía Humana, Facultad de Medicina, Universidad de Castilla-La Mancha, Albacete and the Laboratorio Cajal de Circuitos Corticales UPM-CSIC, Madrid, Spain) (Table S31). Brain tissue was analyzed for Braak stage (*49*) and CERAD neuropathological diagnosis (*50*) and assigned a zero score. Nevertheless, case AB1 showed sparse tau-immunoreactive cells in the hippocampal formation and case AB4 showed a relatively high number of amyloid plaques mainly located in the subicular and the parahippocampal regions. The sampling procedure was approved by the Institutional Ethical Committee. Tissue from some of these human brains has been used in previous unrelated studies (*18, 21*).

After extraction, brain tissue was fixed in cold 4% paraformaldehyde (Sigma-Aldrich, St Louis, MO, USA) in 0.1 M sodium phosphate buffer (PB; Panreac, 131965, Spain), pH 7.4, for 24 h. Subsequently, the block of tissue containing the hippocampus was washed in PB and coronal 150 μm-sections were obtained with a vibratome (Vibratome Sectioning System, VT1200S Vibratome, Leica Biosystems, Germany).

### Tissue processing for EM

Coronal sections from the hippocampal body (*19*) containing the CA1 region were selected and postfixed for 48 h in a solution of 2% paraformaldehyde, 2.5% glutaraldehyde (TAAB, G002, UK) and 0.003% CaCl_2_ (Sigma, C-2661-500G, Germany) in 0.1 M sodium cacodylate buffer (Sigma, C0250-500G, Germany). The sections were treated with 1% OsO_4_ (Sigma, O5500, Germany), 0.1% ferrocyanide potassium (Probus, 23345, Spain) and 0.003% CaCl_2_ in sodium cacodylate buffer (0.1 M) for 1h at room temperature. Sections were then stained with 1% uranyl acetate (EMS, 8473, USA), dehydrated, and flat embedded in Araldite (TAAB, E021, UK) for 48 h at 60°C (*43*). Embedded sections were glued onto a blank Araldite block and trimmed. Semithin sections (1 μm) were obtained from the surface of the block and stained with 1% toluidine blue (Merck, 115930, Germany) in 1% sodium borate (Panreac, 141644, Spain).

### Layer delimitation

The exact location of all CA1 layers was determined by examining 1% toluidine blue-stained semithin sections under a light microscope (Fig. 1). More specifically, the medial portion of the CA1 region was analyzed. From its deepest level to the surface (i.e., from the ventricular cavity towards the vestigial hippocampal sulcus), the cornu ammonis may be divided into five layers: the alveus, *stratum oriens* (SO), *stratum pyramidale* (SP), *stratum radiatum* (SR) and *stratum lacunosum-moleculare* (SLM) (*19*). Within the SP, two sublayers were defined by dividing the layer into a deeper part (dSP; close to the ventricular cavity) and a more superficial part (sSP; close to the vestigial hippocampal sulcus; Fig. 1a-b; Fig. S2) (*51, 52*).

To calculate the thickness of the layers, they were delimited using toluidine blue-stained semithin section adjacent to the block surface (Fig. 1). Three measures per case were taken at different medio-lateral levels of CA1. This analysis was performed using ImageJ (ImageJ 1.51; NIH, USA; http://imagej.nih.gov/ij/).

### Volume fraction estimation of cortical elements

From each case, three semithin sections (1 μm thick; stained with 1% toluidine blue) were used to estimate the volume fraction occupied by blood vessels, cell bodies, and neuropil in each layer. This estimation was performed applying the Cavalieri principle (*13*) by point counting using the integrated Stereo Investigator stereological package (Version 8.0, MicroBrightField Inc., VT, USA) attached to an Olympus light microscope (Olympus, Bellerup, Denmark) at 40x magnification (Fig. S3a). A grid whose points had an associated area of 400 µm^2^ was overlaid over each semithin section to determine the V_v_ occupied by different elements: blood vessels, glia, neurons and neuropil. V_v_ occupied by the neuropil was estimated with the following formula: V_v_ neuropil = 100 - (V_v_ blood vessels + V_v_ glia + V_v_ neurons).

### FIB/SEM technology

A 3D EM study of the samples was conducted using combined FIB/SEM technology (Crossbeam 540 electron microscope, Carl Zeiss NTS GmbH, Oberkochen, Germany). The block of araldite containing the sample was introduced into the microscope, where the FIB removed thin layers of material from the tissue surface on a nanometer scale, allowing the images to be obtained in an automated sequential manner with the backscattered electron detector of the SEM. In this way, long series of photographs of a 3D sample were acquired with a quality and resolution similar to those achieved with TEM (*16*) (Fig. 1c–d). This study was conducted in the neuropil —i.e., avoiding the neuronal and glial somata, blood vessels, large dendrites and myelinated axons— where most synaptic contacts take place (*53*).

Image resolution in the xy plane was 5 nm/pixel. Resolution in the z-axis (section thickness) was 20 nm, and image size was 2048 x 1536 pixels. These parameters were optimized to make it possible to obtain a large enough field of view where the different types of synapses can be clearly identified in a reasonable amount of time (12 h per stack of images). The volume per stack ranged from 356 μm^3^ to 727 μm^3^ (225 and 459 images, respectively). Volume measurements were corrected for tissue shrinkage due to EM processing (*17*) with a shrinkage factor obtained by dividing the area of the tissue after processing by the area of the same tissue before processing (p=0.933). A correction in the volume of the stack of images for the presence of fixation artifact (i.e., swollen neuronal or glial processes) was also applied after quantification with Cavalieri principle (*13*). Every FIB/SEM stack was examined, and the volume artifact ranged from 0 to 20% of the stack volume. A total of 75 stacks of images from all layers of the CA1 field were obtained (3 stacks per case and region in the 5 cases, with a total volume studied of 29,322 μm^3^) (Fig. 1b).

### 3D analysis of synapses

#### Classification of synapses and postsynaptic target identification

EspINA software was used for the 3D segmentation and classification of synapses in the 75 stacks of images (*Espina Interactive Neuron Analyzer*, 2.4.1; Madrid, Spain; https://cajalbbp.es/espina/) (*54*). As previously discussed in (*17*), there is a consensus for classifying cortical synapses into AS (or type I) and SS (or type II) synapses. The main characteristic distinguishing these synapses is the prominent or thin post-synaptic density, respectively. Nevertheless, in single sections, the synaptic cleft and the pre- and post-synaptic densities are often blurred if the plane of the section does not pass at right angles to the synaptic junction. Since the software EspINA allows navigation through the stack of images, it was possible to unambiguously identify every synapse as AS or SS based on the thickness of the PSD. Synapses with prominent PSDs are classified as AS, while thin PSDs are classified as SS (Figs. 1e, 3 and 4).

Additionally, based on the postsynaptic targets, synapses were further classified as axospinous synapses (synapses on dendritic spines) and axodendritic synapses (synapses on dendritic shafts). In the case of axospinous synapses, they were further subdivided into axospinous synapses on the head or on the neck of the spine. For axodendritic synapses, dendritic shafts were further classified as spiny (when dendritic spines could be observed emerging from the shaft) or aspiny. Only clearly identifiable postsynaptic elements were quantified (i.e., elements that were unambiguously identified from navigating through the stack of images; Fig. 4a–k).

Finally, synapses were classified —according to the shape of their synaptic junction— into four categories, as described elsewhere (*36*). In short, synapses with a flat, disk-shaped PSD were classified as macular. A second category was established by the presence of an indentation in the perimeter (horseshoe-shaped synapses). Synaptic junctions with one or more holes in the PSD were referred to as perforated. Synaptic junctions with two or more physically discontinuous PSDs were categorized as fragmented (Fig. 2d).

#### Morphological and spatial measurements

The synaptic apposition surface (SAS) and perimeter of each synaptic junction was extracted with EspINA software to study morphological parameters regarding synapses. This software also permits the quantitation of the curvature of the synapses as it adapts to the curvature of the synaptic junction. Specifically, curvature measurements are calculated as 1 minus the ratio between the projected area of the SAS and the area of the SAS (*54*). This measurement would be 0 in a flat SAS and would increase its value to a maximum of 1 as the SAS curvature increases.

The spatial distribution of synapses was determined by performing a Spatial Point Pattern analysis (*14, 15*). The position of centroids of the synapses was compared to the Complete Spatial Randomness (CSR) model, which defines a situation where a point is equally probable to occur at any location within a given volume. For each stack of images, functions G, F and K were calculated (*55*). In addition, the distance of every synapse to its nearest synapse was measured. This study was carried out using Spatstat package and R Project software (*56*).

### Statistical analysis

Statistical analysis of the data was carried out using GraphPad Prism statistical package (Prism 7.00 for Windows, GraphPad Software Inc., USA), SPSS software (IBM SPSS Statistics for Windows, Version 24.0. Armonk, NY: IBM Corp) and R Project software (R 3.5.1; Bell Laboratories, NJ, USA; http://www.R-project.org). Differences in the V_v_ occupied by cortical elements; synaptic density; and morphological and spatial parameters were analyzed performing either a two-sided, one-way analysis of variance (ANOVA), with Tukey post hoc corrections, or Mann-Whitney U (MW) nonparametric test, as appropriate. Frequency distributions were analyzed using Kolmogorov-Smirnov (KS) nonparametric tests. Chi-squared (χ^2^) tests were used for contingency tables. In general, for any contingency table, the expected frequency for a cell in the i^th^ row and the j^th^ column is E_ij_ = T_i_T_j_/T, where T_i_ is the marginal total for the i^th^ row, T_j_ is the marginal total for the j^th^ column, and T is the total number of observations. χ² tests of association were applied to these tables (*57*). The criterion for statistical significance was considered to be met for p<0.05 when the sample size was equal to the number of subjects (i.e., ANOVA and MW tests), and for p<0.001 when the sample size was equal to the number of synapses (i.e., KS and χ^2^ tests), in order to avoid overestimation of the differences due to a very big sample size.

## Acknowledgments

We would like to thank Carmen Álvarez, Miriam Martín and Lorena Valdés for their helpful technical assistance and Nick Guthrie for his excellent text editing. This work was supported by grants from the following entities: Centro de Investigación en Red sobre Enfermedades Neurodegenerativas (CIBERNED, CB06/05/0066, Spain); the Spanish “Ministerio de Ciencia, Innovación y Universidades” (grant PGC2018-094307-B-I00 and the Cajal Blue Brain Project [the Spanish partner of the Blue Brain Project initiative from EPFL, Switzerland]); the European Union’s Horizon 2020 Research and Innovation Programme under grant agreement No. 785907 (Human Brain Project, SGA2), the Alzheimer’s Association (ZEN-15-321663) and the UNED (Plan de Promoción de la Investigación, 2014-040-UNED-POST). MM-C was awarded a research fellowship from the Spanish Ministry of Education, Culture and Sports (contract FPU14/02245).

## Author contributions

JDeF and LB-L oversaw and designed the project. MM-C and LB-L designed and performed experiments. MM-C carried out data analysis. MD-A and LA-N performed experiments and helped interpret results. PR-C conducted experiments. MM-C and LB-L drafted the initial manuscript. All authors read and approved the final manuscript.

## Competing interests

The authors declare that they have no conflict of interest.

## Data and materials availability

Most data is available in the main text or the Supplementary Materials. The datasets used and analyzed during the current study are available from the corresponding author on request.

