## Supplementary material for "Three-dimensional synaptic organization of the human hippocampal CA1 field"

**This PDF file includes:**

Supplementary Results

Figs. S1 to S8

Tables S1 to S31

### Supplementary Results

#### Electron microscopy

##### *Size of the synapses: SAS Curvature*

While synaptic size parameters area and perimeter were highly correlated ($R^2$ = 0.81 for all synapses; $R^2$ = 0.82 for AS; $R^2$ = 0.81 for SS), curvature measurements showed very little association with either area ($R^2$ = 0.05 for all synapses; $R^2$ = 0.05 for AS; $R^2$ = 0.00 for SS) or perimeter ($R^2$ = 0.08 for all synapses; $R^2$ = 0.09 for AS; $R^2$ = 0.00 for SS). Consequently, differences observed in these two parameters did not extrapolate to variations in the curvature (Fig. S8).

No differences in the curvature of the synapses were observed between AS and SS (mean SAS curvature of AS: 0.050; mean SAS curvature of SS: 0.047; Fig. S8; Table S12). Likewise, no curvature differences were seen between the layers (KS, p>0.001; Fig. S8).

The frequency histograms of SAS curvature ratios showed a positive skewness with a greater proportion of synapses presenting lower values, meaning a larger prevalence of flatter synapses than more curved ones for both AS and SS populations, but also within every layer, with great overlap among all the distributions (KS, p>0.001; Fig. S8).

Regarding the postsynaptic targets, the curvature ratio of axospinous and axodendritic synapses did not differ for either population — AS or SS (MW, p>0.05; Fig. S8; Table S13).

When focusing on the shape of the synaptic junction, fragmented AS were found to be more curved than macular AS (ANOVA, p=0.04) — a difference that was maintained through all layers (ANOVA, p<0.001; Fig. S8; Table S14). In the case of SS, fragmented SS were observed to be more curved than the rest of the synaptic shape types in SLM (ANOVA, p=0.0480 for macular SS-fragmented SS; p=0.0050 for HS SS-fragmented SS; and p=0.0009 for perforated

SS-fragmented SS; Fig. S8; Table S14). Differences in the mean SAS curvature were also observed within the same synaptic shape type between layers. In this regard, SLM presented flatter horseshoe-shaped AS than both dSP (ANOVA, p=0.0035) and sSP (p=0.0005), while SO exhibited flatter perforated AS than sSP (ANOVA, $p=1.246\times10^{-5}$) and SR (ANOVA, $p=5.538\times10^{-5}$; Fig. S8; Table S14).

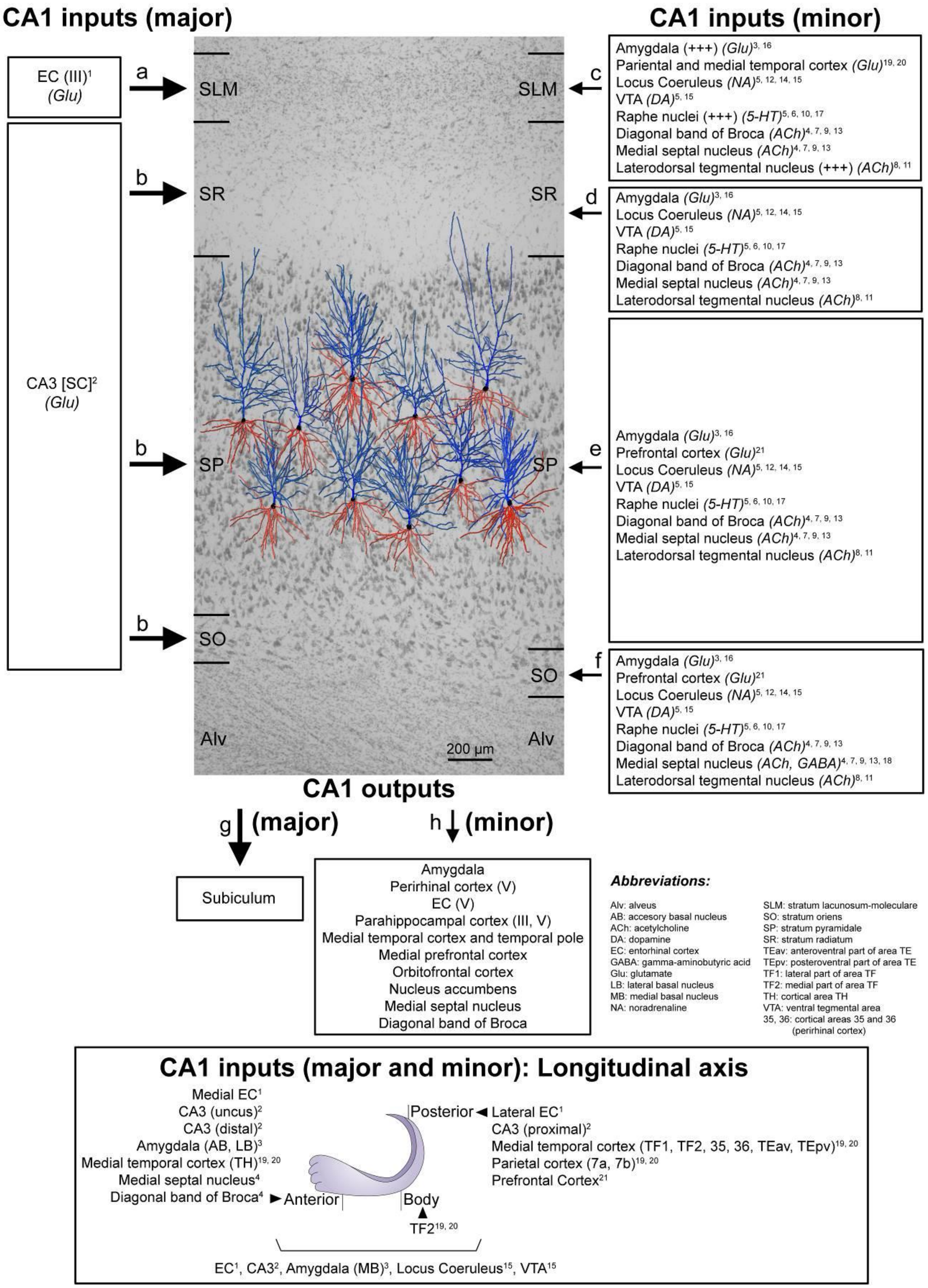

**Fig. S1.** Schematic representation of the main direct connections between CA1 and other brain regions in primates (monkeys, unless otherwise specified; note that all abbreviations are defined in the figure itself). Photomicrograph of a Nissl-stained coronal brain section from the human

CA1 (in black and white) with reconstructed pyramidal neurons (taken from *27*) superimposed on the same scale. The pyramidal neurons have been placed in the middle of the SP, approximately where they were injected with Lucifer Yellow. The apical and basal dendritic arbors are colored in blue and red, respectively. Major and minor projections have been represented with large and small arrows, respectively.

The major input to CA1 originates in the EC (glutamatergic) (a–b). Neurons located in layer III of the EC project directly to SLM (a) (*50*; 1), while layer II neurons project indirectly to SO and SR, but, unlike in rodents, they also project to SP (b), via DG and CA3 Schaffer collaterals (SC) (glutamatergic) (*51;* 2). Additionally, minor projections —from the amygdala (glutamatergic), the VTA (dopaminergic), the locus coeruleus (noradrenergic), the raphe nuclei (serotoninergic), the medial septal nucleus (Ch1) (cholinergic), the vertical limb of the diagonal band of Broca (cholinergic) and the laterodorsal tegmental nucleus (cholinergic)— have been described to arrive in all layers of CA1 (c–f). However, some of these projections (amygdala, raphe nuclei and laterodorsal tegmental nucleus) have been described as being especially numerous in the SLM (c, represented with a "+++") (*52*; 3); (*53*; 4); (*54*; 5); (*55*; 6); (*56*; 7); (*57*; 8); (*58*; 9); (*59*; 10); (*60*; 11); (*61*; 12); (*62*; 13); (*63*; 14); (*64*; 15); (*65*; 16); (*66*; 17). In addition, differences have been reported regarding the density of immunoreactive ChAT fibers between human and monkey (higher density of fiber labeling in SP and SR in human CA1 and higher density in SLM and SO in monkey CA1) (*53*; 4), (*56*; 7). Minor direct projections have also been reported from the medial septal nuclei (GABAergic) to SO (mainly innervating interneurons) (f) (*67*; 18). In addition, minor projections have also been reported from several cortical regions (glutamatergic) including: the medial temporal cortex (TH, TF1, TF2 — posterior parahippocampal areas; 35, 36 — perirhinal cortex; TEav, TEpv —ventral inferotemporal areas) and the parietal cortex (7a and 7b) mainly to SLM (c) (*68*; 19) (*69*; 20) and from the prefrontal cortex mainly to SO and SP (e,f) (*70*; 21). Other regions have been observed to project to the hippocampal formation of the monkey but there is no specific information regarding possible direct projections to CA1 (for example, the claustrum; the substantia innominate; the basal nucleus of Meynert; the thalamus (specifically the anterior nuclear complex, the laterodorsal nucleus, the paraventricular and parataenial nuclei, the nucleus reuniens, and the nucleus centralis medialis); the lateral preoptic and lateral hypothalamic areas; the supramammillary and retromammillary regions; the tegmental reticular fields; the nucleus reticularis tegmenti pontis; and the central gray — for more details, see *54* (4). For example, the midline thalamic nuclei (mostly Reuniens nucleus) have been observed to project mainly to the SLM in the rat CA1 but there is no information regarding primates (see *71*).

Regarding the longitudinal axis of the hippocampus (bottom of the diagram modified from *72*), there are some examples in which it has been described that primate CA1 receives different inputs along this axis. For example, (*i*) lateral portions of the EC project to caudal levels of the recipient fields and more medial parts of the EC project to progressively more rostral portions (*50*; 1); (*ii*) CA3 projections to CA1 extend very widely both rostrally and caudally. However, the projections from CA3 neurons located at the level of the uncus are restricted to rostral CA1,

whereas neurons in proximal CA3 project to caudal CA1 and neurons in distal CA3 project to rostral CA1 (*51*; 2); (*iii*) the accessory and lateral basal nucleus of the amygdala project to rostral CA1, while the medial basal nucleus projects along the whole longitudinal axis (*52*; 3); (*iv*) the temporal TH cortex projects to rostral CA1, the temporal TF2 cortex to medial CA1 and temporal TF1, TF2, 35, 36, TEav and TEpv and the parietal (7a and 7b) cortices project to caudal CA1 (*68*; 19); (*69*; 20); (*v*) the cholinergic innervation from medial septal nuclei and diagonal band of Broca is mainly present in rostral CA1 (*53*; 4); (see also *73*).
The main output of the CA1 region is the subiculum (i; *71*). However, CA1 has also been reported to project directly to other brain areas (j), such as the amygdala, the orbitofrontal cortex (areas 11 and 13), the medial prefrontal cortex (areas 14, 25 and 32), the medial temporal area TE and the temporal pole TG, to a similar degree as the subicular projection (*52,73-80*). Furthermore, CA1 projects to other areas including layer V of both the EC and the perirhinal cortex, as well as layers III and V of the parahippocampal cortex, where CA1 becomes the major hippocampal source of projections (*69,73,77,81,82*). Additionally, CA1 sends projections to the medial septal nucleus (Ch1), vertical limb of the diagonal band of Broca (Ch2) and the nucleus accumbens, and it is known that these projections are not shared with the subiculum (*73,80,83,84*). According to *85*, CA1 does not project to the thalamus.
Finally, there are also differences in the hippocampal projections along the longitudinal hippocampal axis (*72,73,86*), For example, more rostral parts of CA1 project mainly to rostral and medial parts of the perirhinal cortex, as well as to the lateral septum, the amygdala, nucleus accumbens, the orbitofrontal cortex and the medial prefrontal cortex (*52,73-76,78*). Mid portions of CA1 project to the EC, while the most posterior parts of CA1 send projections mainly to the caudal and lateral portions of the parahippocampal cortex and, to a lesser extent, to dorsal and medial parts of the septum (*69,73,77,87*). SLM: *stratum lacunosum-moleculare*; SO: *stratum oriens*; SP: *stratum pyramidale*, SR: *stratum radiatum*.

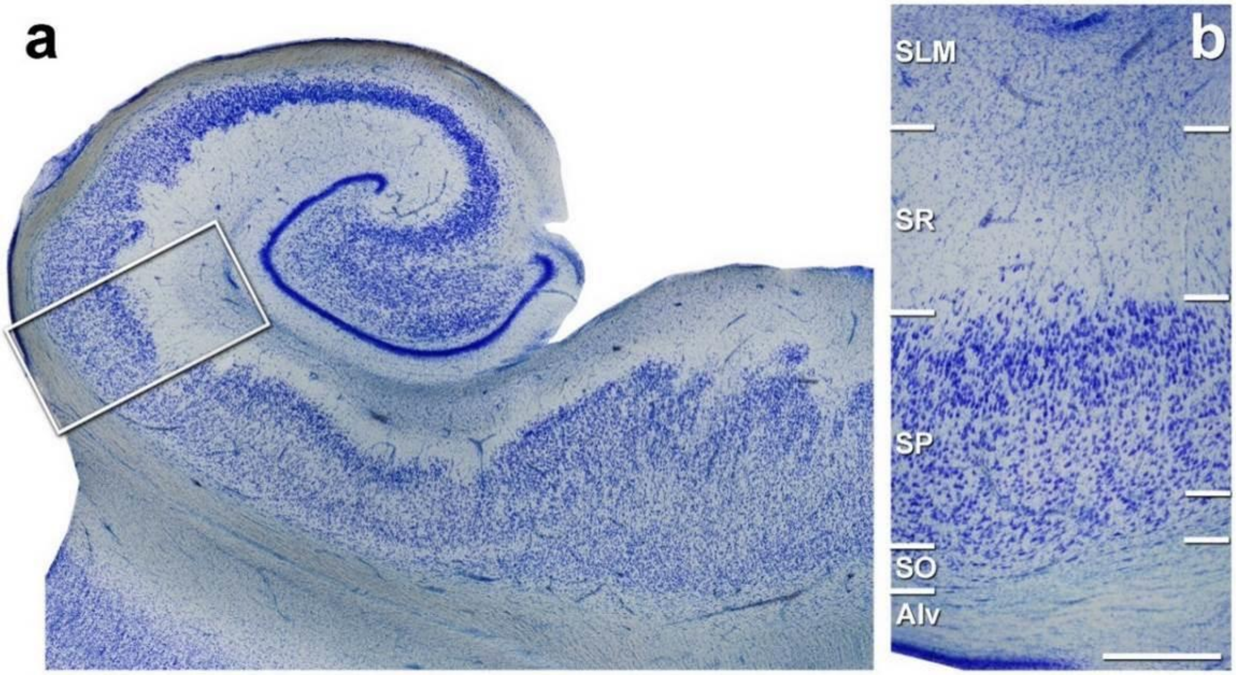

**Fig. S2.** Coronal section of the human hippocampus at the level of the hippocampal body. **a**, Low-power photomicrograph of a Nissl-stained coronal brain section from human hippocampus. **b**, Higher magnification image of the boxed area shown in (**a**) showing the medial CA1 hippocampal field. Alv: alveus; SO, *stratum oriens*: SP, *stratum pyramidale*; SR: *stratum radiatum*; SLM: *stratum lacunosum-moleculare*. Scale bar in b corresponds to: 1430 µm in a, 490 µm in b.

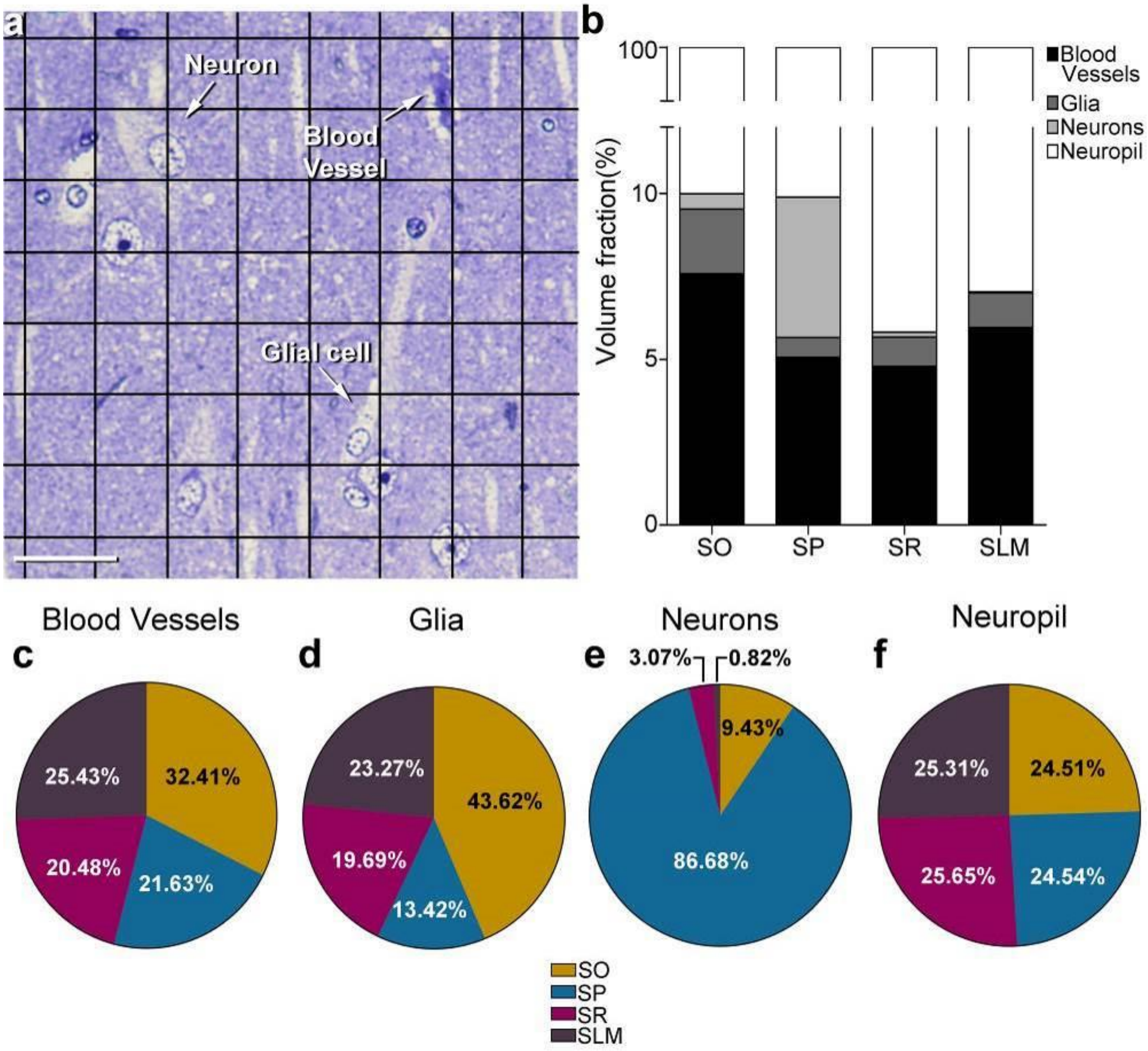

**Fig. S3.** Stereological estimation of the volume occupied by different cortical elements in the CA1 using a stereological grid. **a**, photomicrograph of a toluidine blue-stained 1 µm thick semithin section. Points hitting the different cortical elements (blood vessels, soma of glial cells and neurons) are labelled and indicated by arrows. **b**, the volume fraction occupied by each cortical element in every stratum is represented with a bar chart. **c-f**, shown by pie charts the relative volume fraction including all layers of blood vessels (**c**), glial cells (**d**), neurons (**e**) and neuropil (**f**). dSP: deep part of *stratum pyramidale*; SLM: *stratum lacunosum-moleculare*; SO: *stratum oriens*; SR: *stratum radiatum*; sSP: superficial part of *stratum pyramidale*. Scale bar in a: 30 µm.

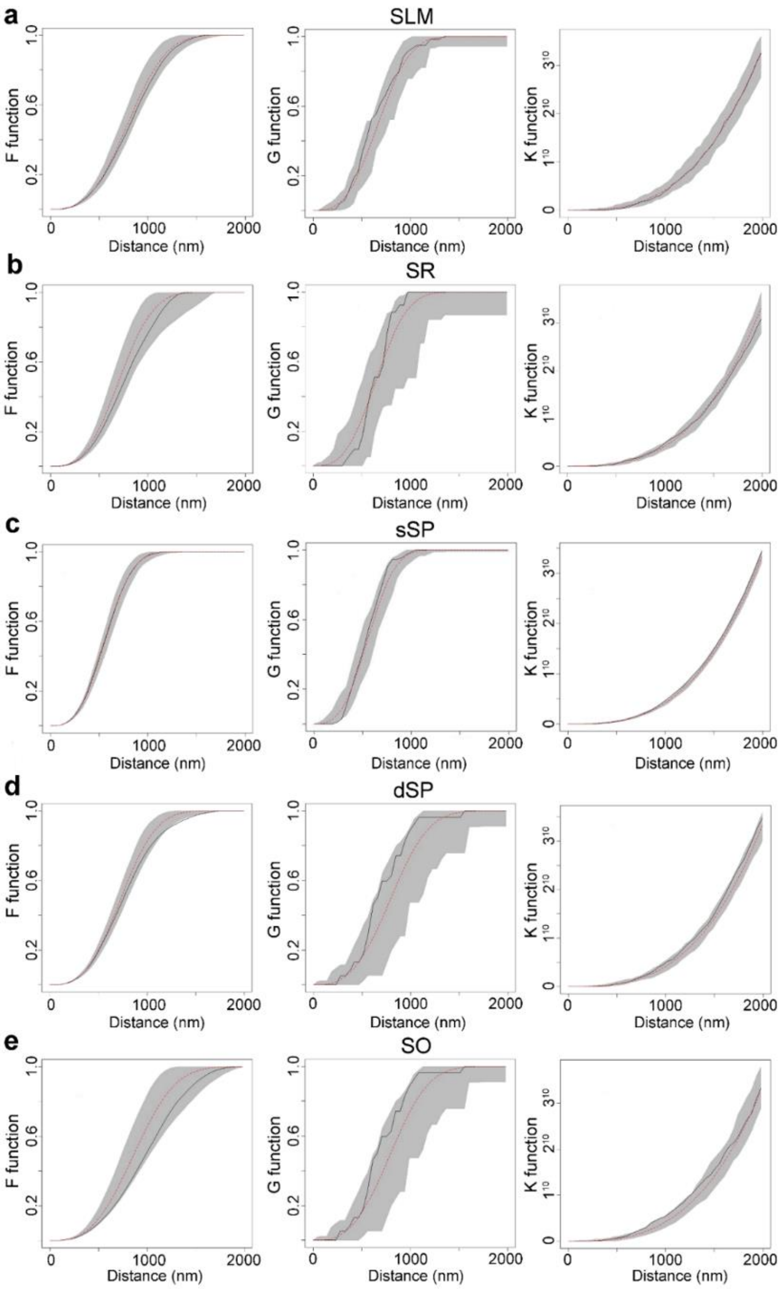

**Fig. S4.** Analysis of the synaptic spatial distribution in the neuropil. **a–e**, F, G and K functions obtained in each layer: SLM (**a**), SR (**b**), sSP (**c**), dSP (**d**) and SO (**e**). Red dashed traces correspond to a theoretical homogeneous Poisson process for each function (F, G, K). The black continuous traces correspond to the experimentally observed function. The shaded areas represent the envelopes of values calculated from a set of 99 simulations. All plots show a distribution which fits a Poisson function in all strata which is representative of a random spatial distribution. SO, *stratum oriens*; SP, *stratum pyramidale*; SR, *stratum radiatum*; SLM, *stratum lacunosum-moleculare*.

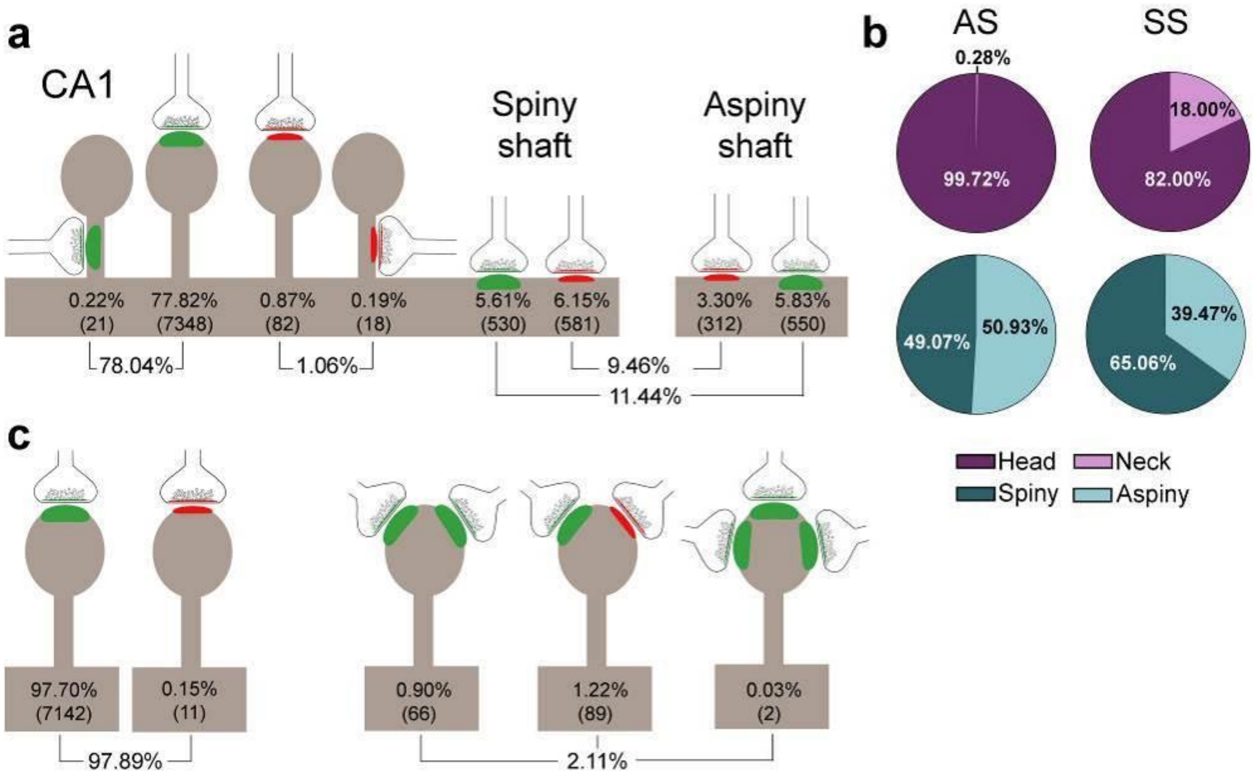

**Fig. S5.** Schematic representation of the distribution of synapses regarding postsynaptic targets and dendritic spines in the whole CA1. **a**, Frequency of axospinous synapses (both on the head and neck of dendritic spines) and axodendritic synapses (both on spiny and aspiny shafts) for asymmetric synapses (AS; green) and symmetric synapses (SS; red). Percentage and number of each synaptic type (below in brackets) are shown. **b**, Proportions of the exact location of axospinous synapses (i.e., the head or neck of the spine) and axodendritic synapses (i.e., spiny or aspiny shafts) in both AS and SS, represented as pie charts. **c**, Percentage of dendritic spines receiving single or multiple synaptic contacts.

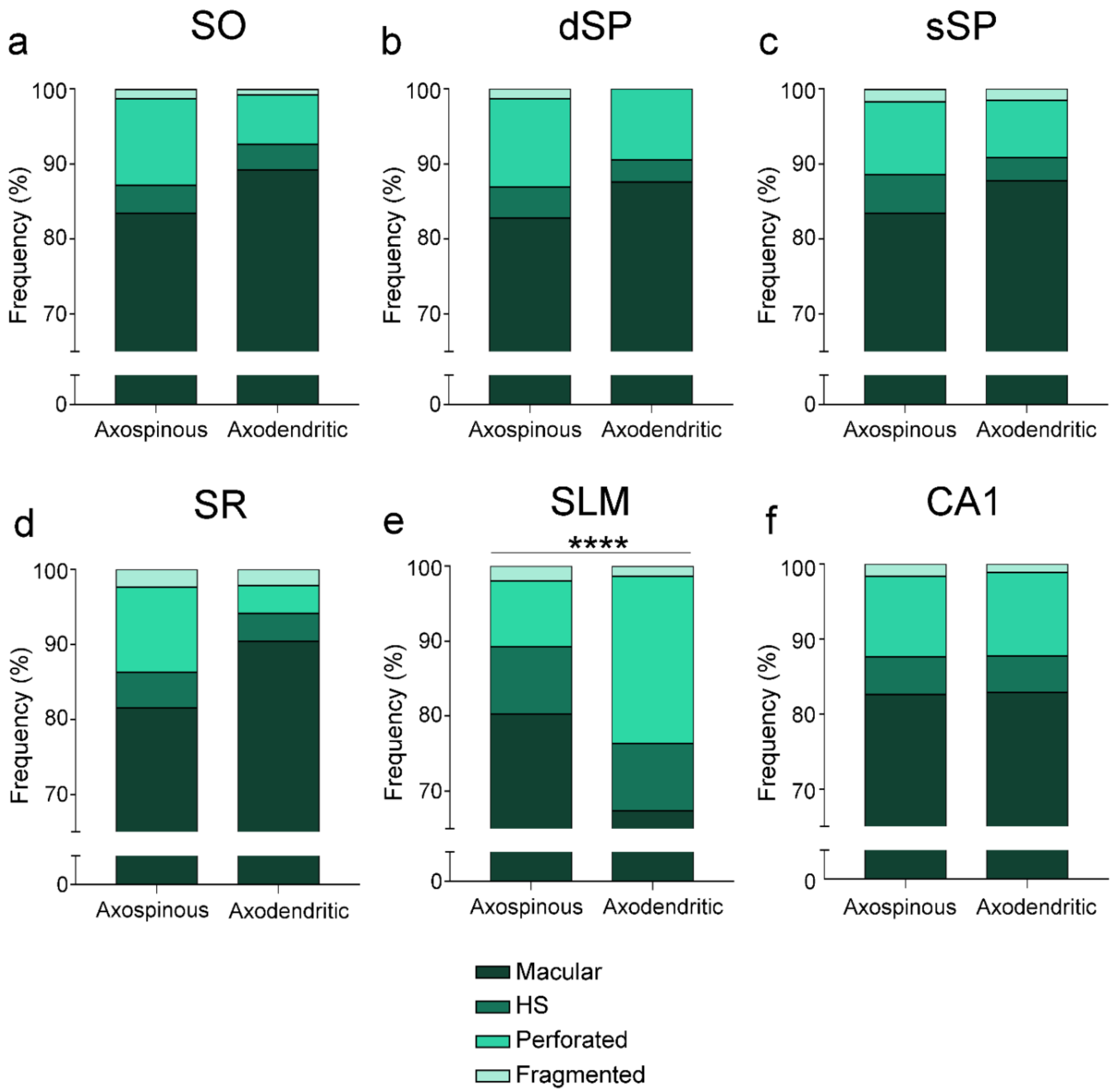

**Fig. S6.** Frequency plots for every type of synaptic shape found among axospinous AS and axodendritic AS. **a–f**, Plots are shown for all layers (**a–e**) and for the CA1 region as a whole (**f**). Only in SLM, non-macular synapses are more abundant in the axodendritic AS population than in the axospinous AS one. Moreover, overall, SLM presents a higher frequency of complex-shaped synapses than other layers (see Figure 2). AS: asymmetric synapses; SO, *stratum oriens*; dSP, deep *stratum pyramidale*; sSP, superficial *stratum pyramidale*; SR, *stratum radiatum*; SLM, *stratum lacunosum-moleculare*.

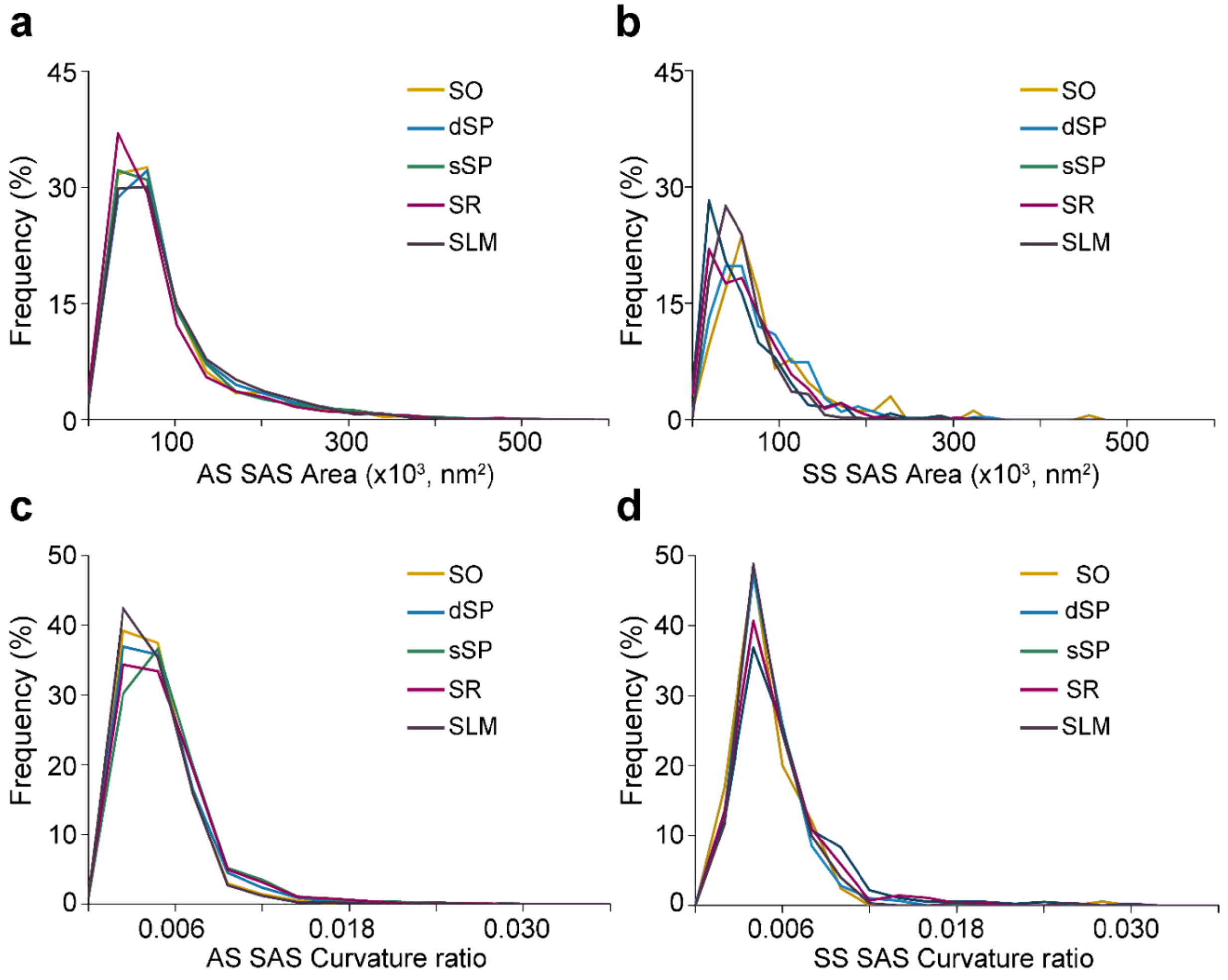

**Fig. S7.** Frequency distribution histograms of the synaptic surface area (SAS) and curvature in CA1. **a,b**, Frequency distribution histograms of SAS area in all layers of CA1 for asymmetric synapses (AS) (**a**) and symmetric synapses (SS) (**b**). Note there is a clear overlap in the frequency distribution among layers both for AS and SS. **c,d,** Frequency distribution histograms of SAS curvature ratio in every CA1 layer for AS (**c**) and SS (**d**). Note there is a clear overlap in the frequency distribution among layers both for AS and SS. All histograms show a positive skewness with a higher frequency of smaller values than bigger ones. SO, *stratum oriens*; dSP, deep *stratum pyramidale*; sSP, superficial stratum pyramidale; SR, stratum radiatum; SLM, stratum lacunosum-moleculare.

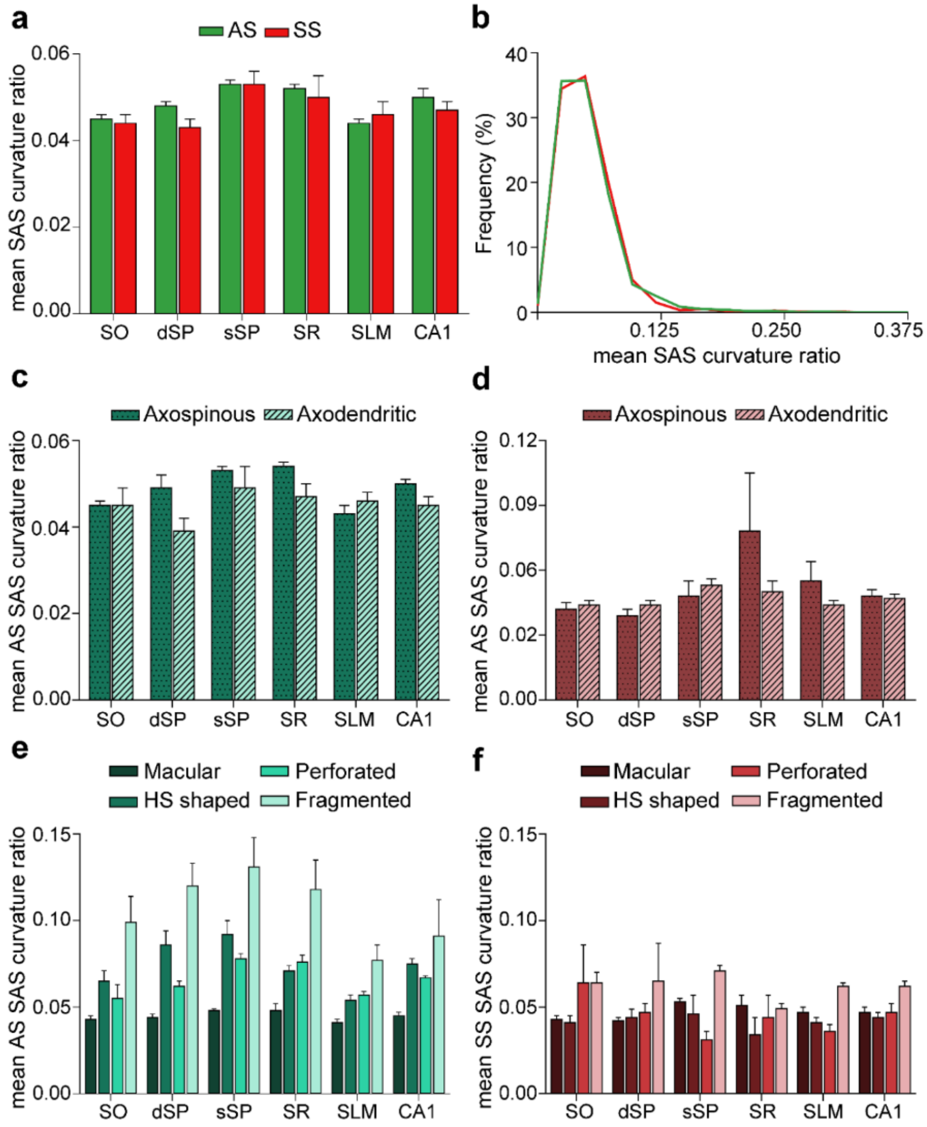

**Fig. S8.** Synaptic size: synaptic surface area (SAS) curvature ratio. **a**, Mean SAS curvature ratio of asymmetric synapses (AS) and symmetric synapses (SS) are represented for each layer and all layers of CA1. No differences were observed in the curvature ratio between AS and SS. **b**, Frequency distribution of the SAS curvature ratio for AS and SS in all layers. Histograms show a positive skewness with a higher frequency of smaller values than bigger ones, both for AS and SS. **c,d,** The mean SAS curvature of axospinous and axodendritic synapses has been plotted for AS (**c**) and for SS (**d**) in each layer and all layers. No differences were observed for either AS (**c**) or SS (**d**). **e–f,** The mean SAS curvature of synapses according to their shape has been plotted for AS (**e**) and for SS (**f**) in each layer and all layers. Differences in the curvature ratio between the different synaptic shapes were observed. Fragmented AS were more curved than macular AS in the whole CA1 (**e**) — a difference that was maintained throughout all layers (ANOVA, ranged from $p<0.001$ to $p<0.0001$). Additionally, fragmented SS were more curved than the rest of the morphological types of SS (**f**), although this difference was only significant in SLM when taking the layers into account (ANOVA, $p<0.05$). Differences in the mean SAS curvature could also be observed within the same synaptic shape type between layers. In this regard, SLM presented flatter HS AS than both dSP (ANOVA, $p<0.01$) and sSP ($p<0.001$), while SO exhibited flatter perforated AS than sSP and SR (ANOVA, $p<0.0001$). SO, *stratum oriens*; dSP, deep *stratum pyramidale*; sSP, superficial *stratum pyramidale*; SR, *stratum radiatum*; SLM, *stratum lacunosum-moleculare*.

| | Thickness (mm; mean±SD) | Percentage of all CA1 layers | $V_{bv}$ (%; mean±SD) | $V_g$ (%; mean±SD) | $V_{neu}$ (%; mean±SD) | $V_n$ (%; mean±SD) |
|---|---|---|---|---|---|---|
| SO | 0.06±0.03 | 2% | 7.58±3.29 | 1.95±0.83 | 0.46±0.47 | 90.01±3.07 |
| SP | 1.13±0.33 | 42% | 5.06±0.99 | 0.60±0.30 | 4.23±1.07 | 90.11±1.32 |
| SR | 0.55±0.31 | 20% | 4.79±0.75 | 0.88±0.75 | 0.15±0.20 | 94.19±1.17 |
| SLM | 0.62±0.16 | 23% | 5.96±1.25 | 1.04±0.78 | 0.04±0.03 | 92.95±0.73 |
| All layers | 2.70±0.62 | 100% | - | - | - | - |

**Table S1.**

Data on CA1 thickness and volume fraction occupied by different cortical elements per layer. SD: standard deviation; SLM: *stratum lacunosum-moleculare*; SO: *stratum oriens*; SP: *stratum pyramidale*; SR: *stratum radiatum*; $V_{bv}$: volume fraction occupied by blood vessels; $V_g$: volume fraction occupied by glia; $V_{neu}$: volume fraction occupied by neurons; $V_n$: volume fraction occupied by neuropil.

| | SO | dSP | sSP | SR | SLM | Totals |
|---|---|---|---|---|---|---|
| Axospinous AS | **1359 (76.48%)** | **1788 (81.16%)** | **2184 (87.61%)** | **1278 (78.36%)** | **760 (56.80%)** | **7369 (78.04%)** |
| *On the head* | 1354 | 1784 | 2178 | 1273 | 759 | 7348 |
| *On the neck* | 5 | 4 | 6 | 5 | 1 | 21 |
| Axospinous SS | **9 (0.51%)** | **37 (1.68%)** | **17 (0.68%)** | **6 (0.37%)** | **31 (2.32%)** | **100 (1.06%)** |
| *On the head* | 7 | 30 | 11 | 5 | 29 | 82 |
| *On the neck* | 2 | 7 | 6 | 1 | 2 | 18 |
| Axodendritic AS | **259 (14.58%)** | **202 (9.17%)** | **131 (5.25%)** | **188 (11.53%)** | **300 (22.42%)** | **1080 (11.44%)** |
| *On spiny shafts* | 165 | 107 | 82 | 95 | 81 | 530 |
| *On aspiny shafts* | 94 | 95 | 49 | 93 | 219 | 550 |
| Axodendritic SS | **150 (8.44%)** | **176 (7.99%)** | **161 (6.46%)** | **159 (9.75%)** | **247 (18.46%)** | **893 (9.46%)** |
| *On spiny shafts* | 98 | 144 | 121 | 114 | 104 | 581 |
| *On aspiny shafts* | 52 | 32 | 40 | 45 | 143 | 312 |
| Axospinous AS+SS | **1368 (76.98%)** | **1825 (82.84%)** | **2201 (88.29%)** | **1284 (78.72%)** | **791 (59.12%)** | **7469 (79.10%)** |
| *On the head* | 1361 (76.59%) | 1814 (82.34%) | 2189 (87.81%) | 1278 (78.36%) | 788 (58.89%) | 7430 (78.69%) |
| *On the neck* | 7 (0.39%) | 11 (0.50%) | 12 (0.48%) | 6 (0.37%) | 3 (0.22%) | 39 (0.41%) |
| Axodendritic AS+SS | **409 (23.02%)** | **378 (17.16%)** | **292 (11.71%)** | **347 (21.28%)** | **547 (40.88%)** | **1973 (20.90%)** |
| *On spiny shafts* | 263 (14.80%) | 251 (11.39%) | 203 (8.14%) | 209 (12.81%) | 185 (13.83%) | 1111 (11.77%) |
| *On aspiny shafts* | 146 (8.22%) | 127 (5.76%) | 89 (3.57%) | 138 (8.46%) | 362 (27.06%) | 862 (9.13%) |
| Total AS+SS | **1777 (100%)** | **2203 (100%)** | **2493 (100%)** | **1631 (100%)** | **1338 (100%)** | **9442 (100%)** |

**Table S2.**

Postsynaptic target information in all layers of CA1. More detailed information on axospinous synapses can be found in Table S9. AS: asymmetric synapses; dSP: deep part of *stratum pyramidale*; SLM: *stratum lacunosum-moleculare*; SO: *stratum oriens*; SR: *stratum radiatum*; SS: symmetric synapses; sSP: superficial part of *stratum pyramidale.*

| | Type of postsynaptic target | | | | |
|---|---|---|---|---|---|
| | Axospinous AS | Axodendritic AS | Axodendritic SS | Axospinous SS | Totals |
| SO | **1359** (1386.86) | **259** (203.26) | **150** (168.06) | **9** (18.82) | 1777 |
| dSP | **1788** (1719.33) | **202** (251.98) | **176** (208.35) | **37** (23.33) | 2203 |
| sSP | **2184** (1945.66) | **131** (285.16) | **161** (235.78) | **17** (26.40) | 2493 |
| SR | **1278** (1272.91) | **188** (186.56) | **159** (154.26) | **6** (17.27) | 1631 |
| SLM | **760** (1044.24) | **300** (153.04) | **247** (126.54) | **31** (14.17) | 1338 |
| Totals | 7369 | 1080 | 893 | 100 | 9442 |

**Table S3.**

A 5x4 contingency table generated to compare the proportion of synapses according to their postsynaptic targets between all layers of CA1. The observed counts of synapses in each subcategory are shown in bold. The expected counts (in parentheses) are calculated from the marginal totals, assuming the null hypothesis that there is no association between the layer and the type of postsynaptic target. A $\chi^2$ test of association was applied to the table, indicating that the null hypothesis must be rejected ($p<0.0001$). Further 2x2 contingency tables (40 in total; some examples are shown in Tables S5–S8) were created in order to find the exact layers and postsynaptic targets presenting differences in prevalence, comparing each layer to one another and to every type of postsynaptic target. Similar tables were created to compare all CA1 layers in the rest of the categories analyzed (e.g., excitatory vs inhibitory contacts, synapse subtypes based on the shape of their synaptic junction, etc.).

| | Type of postsynaptic target | | |
|---|---|---|---|
| | Axospinous AS | Other synaptic types | Totals |
| SO | **1359** (1405.08) | **418** (371.92) | 1777 |
| dSP | **1788** (1741.92) | **415** (461.08) | 2203 |
| Totals | 3147 | 833 | 3980 |

**Table S4.**

2x2 contingency table generated after the 5x4 contingency table analysis to compare the proportion of axospinous AS between layers SO and dSP. A $\chi^2$ test of association was applied to the table, indicating that the null hypothesis must be rejected (p<0.001).

| | Type of postsynaptic target | | |
|---|---|---|---|
| | Axodendritic AS | Other synaptic types | Totals |
| SO | **259** (205.83) | **1518** (1571.17) | 1777 |
| dSP | **202** (255.17) | **2001** (1947.83) | 2203 |
| Totals | 461 | 3519 | 3980 |

**Table S5.**

2x2 contingency table generated after the 5x4 contingency table analysis to compare the proportion of axodendritic AS between layers SO and dSP. A $\chi^2$ test of association was applied to the table, indicating that the null hypothesis must be rejected ($p<0.0001$).

| | Type of postsynaptic target | | |
|---|---|---|---|
| | Axodendritic SS | Other synaptic types | Totals |
| SO | **150** (145.55) | **1627** (1631.45) | 1618 |
| dSP | **176** (180.45) | **2027** (2022.55) | 1466 |
| Totals | 326 | 3654 | 3980 |

**Table S6.**

2x2 contingency table generated after the 5x4 contingency table analysis to compare the proportion of axodendritic SS between layers SO and dSP. A $\chi^2$ test of association was applied to the table, indicating that the null hypothesis must be accepted ($p>0.05$).

| | Type of postsynaptic target | | |
|---|---|---|---|
| | Axospinous SS | Other synaptic types | Totals |
| SO | **9** (19.05) | **1768** (1755.95) | 1775 |
| dSP | **37** (24.95) | **2287** (2299.05) | 2324 |
| Totals | 44 | 4055 | 4099 |

**Table S7.**

2x2 contingency table generated after the 5x4 contingency table analysis to compare the proportion of axospinous SS between layers SO and dSP. A $\chi^2$ test of association was applied to the table, indicating that the null hypothesis must be rejected ($p<0.001$).

| | | SO | dSP | sSP | SR | SLM | Totals |
|---|---|---|---|---|---|---|---|
| AS population | Axospinous AS | 1359 (83.99%) | 1788 (89.85%) | 2184 (94.34%) | 1278 (87.18%) | 760 (71.70%) | 7369 (87.22%) |
| | Axodendritic AS | 259 (16.01%) | 202 (10.15%) | 131 (5.66%) | 188 (12.82%) | 300 (28.30%) | 1080 (12.78%) |
| | Total | 1618 (100%) | 1990 (100%) | 2315 (100%) | 1466 (100%) | 1060 (100%) | 8449 (100%) |
| SS population | Axospinous SS | 9 (5.66%) | 37 (17.37%) | 17 (9.55%) | 6 (3.64%) | 31 (11.15%) | 100 (10.07%) |
| | Axodendritic SS | 150 (94.34%) | 176 (82.63%) | 161 (90.45%) | 159 (96.36%) | 247 (88.85%) | 893 (89.93%) |
| | Total | 159 (100%) | 213 (100%) | 178 (100%) | 165 (100%) | 278 (100%) | 993 (100%) |
| Axospinous synapses | AS | 1359 (99.34%) | 1788 (97.97%) | 2184 (99.23%) | 1278 (99.53%) | 760 (96.08%) | 7369 (98.66%) |
| | SS | 9 (0.66%) | 37 (2.03%) | 17 (0.77%) | 6 (0.47%) | 31 (3.92%) | 100 (1.34%) |
| | Total | 1368 (100%) | 1825 (100%) | 2201 (100%) | 1284 (100%) | 791 (100%) | 7469 (100%) |
| Axodendritic synapses | AS | 259 (63.33%) | 202 (53.44%) | 131 (44.86%) | 188 (54.18%) | 300 (54.84%) | 1080 (54.74%) |
| | SS | 150 (36.67%) | 176 (46.56%) | 161 (55.14%) | 159 (45.82%) | 247 (45.16%) | 893 (45.26%) |
| | Total | 409 (100%) | 378 (100%) | 292 (100%) | 347 (100%) | 547 (100%) | 1973 (100%) |

**Table S8.**

Postsynaptic target information of AS and SS in all layers of CA1. More detailed information on axospinous synapses can be found in Table S9. AS: asymmetric synapses; dSP: deep part of *stratum pyramidale*; SLM: *stratum lacunosum-moleculare*; SO: *stratum oriens*; SR: *stratum radiatum*; SS: symmetric synapses; sSP: superficial part of *stratum pyramidale.*

| | SO | dSP | sSP | SR | SLM | CA1 |
|---|---|---|---|---|---|---|
| Spines with one synapse | 1338 (98.89%) | 1740 (97.64%) | 2118 (98.10%) | 1240 (98.26%) | 717 (95.09%) | 7153 (97.89%) |
| 1 AS | 1338 | 1733 | 2118 | 1240 | 713 | 7142 |
| 1 SS | 0 | 7 | 0 | 0 | 4 | 11 |
| Spines with multiple synapses | 15 (1.11%) | 42 (2.36%) | 41 (1.90%) | 22 (1.74%) | 37 (4.91%) | 157 (2.11%) |
| 2 AS | 6 | 11 | 23 | 16 | 10 | 66 |
| 1 AS+1 SS | 9 | 30 | 17 | 6 | 27 | 89 |
| 3 AS | 0 | 1 | 1 | 0 | 0 | 2 |
| Spines with 1 head | 1327 (99.03%) | 1743 (98.03%) | 2117 (98.37%) | 1246 (98.75%) | 752 (99.75%) | 7185 (98.61%) |
| Spines with multiple heads | 13 (0.97%) | 35 (1.97%) | 35 (1.63%) | 16 (1.25%) | 2 (0.25%) | 101 (1.39%) |
| 2 Heads | 12 | 34 | 33 | 16 | 2 | 97 |
| 3 Heads | 1 | 1 | 2 | 0 | 0 | 4 |

**Table S9.**

Data on dendritic spines in each layer of CA1. AS: asymmetric synapses; CA: *cornu ammonis*; dSP: deep part of *stratum pyramidale*; SLM: *stratum lacunosum-moleculare*; SO: *stratum oriens*; SR: *stratum radiatum*; SS: symmetric synapses; sSP: superficial part of *stratum pyramidale.*

| | Type of synapse | No. macular synapses | No. horseshoe-shaped synapses | No. perforated synapses | No. fragmented synapses |
|---|---|---|---|---|---|
| SO | AS | 2280 (86.10%) | 86 (3.25%) | 250 (9.44%) | 32 (1.21%) |
| | SS | 133 (80.12%) | 22 (13.25%) | 8 (4.82%) | 3 (1.81%) |
| dSP | AS | 3294 (85.58%) | 141 (3.66%) | 357 (9.28%) | 57 (1.48%) |
| | SS | 215 (76.51%) | 39 (13.88%) | 24 (8.54%) | 3 (1.07%) |
| sSP | AS | 4517 (87.15%) | 212 (4.09%) | 371 (7.16%) | 83 (1.60%) |
| | SS | 164 (83.67%) | 16 (8.16%) | 6 (3.06%) | 10 (5.10%) |
| SR | AS | 3338 (87.02%) | 150 (3.91%) | 274 (7.14%) | 74 (1.93%) |
| | SS | 141 (81.98%) | 21 (12.21%) | 3 (1.74%) | 7 (4.07%) |
| SLM | AS | 2160 (82.38%) | 189 (7.21%) | 222 (8.47%) | 51 (1.95%) |
| | SS | 258 (81.65%) | 37 (11.71%) | 13 (4.11%) | 8 (2.53%) |
| Totals | AS | 15589 (85.95%) | 778 (4.29%) | 1474 (8.13%) | 297 (1.64%) |
| | SS | 911 (80.55%) | 135 (11.94%) | 54 (4.77%) | 31 (2.74%) |

**Table S10.**

Proportion of AS and SS according to the shape of their synaptic junction in every layer of CA1. AS: asymmetric synapses; dSP: deep part of *stratum pyramidale*; SLM: *stratum lacunosum-moleculare*; SO: *stratum oriens*; SR: *stratum radiatum*; SS: symmetric synapses; sSP: superficial part of *stratum pyramidale.*

| | Type of synapse | No. macular synapses | No. horseshoe-shaped synapses | No. perforated synapses | No. fragmented synapses |
|---|---|---|---|---|---|
| SO | Axospinous AS | 1134 (83.44%) | 51 (3.75%) | 156 (11.48%) | 18 (1.32%) |
| | Axodendritic AS | 231 (89.19%) | 9 (3.47%) | 17 (6.56%) | 2 (0.77%) |
| dSP | Axospinous AS | 1480 (82.77%) | 74 (4.14%) | 211 (11.80%) | 23 (1.29%) |
| | Axodendritic AS | 177 (87.62%) | 6 (2.97%) | 19 (9.41%) | 0 (0.00%) |
| sSP | Axospinous AS | 1822 (83.42%) | 113 (5.17%) | 213 (9.75%) | 36 (1.65%) |
| | Axodendritic AS | 115 (87.79%) | 4 (3.05%) | 10 (7.63%) | 2 (1.53%) |
| SR | Axospinous AS | 1042 (81.53%) | 61 (4.77%) | 145 (11.35%) | 30 (2.35%) |
| | Axodendritic AS | 170 (90.43%) | 7 (3.72%) | 7 (3.72%) | 4 (2.13%) |
| SLM | Axospinous AS | 610 (80.26%) | 68 (8.95%) | 67 (8.82%) | 15 (1.97%) |
| | Axodendritic AS | 202 (67.33%) | 27 (9.00%) | 67 (22.33%) | 4 (1.33%) |
| Totals | Axospinous AS | 6088 (82.62%) | 367 (4.98%) | 792 (10.75%) | 122 (1.66%) |
| | Axodendritic AS | 895 (82.87%) | 53 (4.91%) | 120 (11.11%) | 12 (1.11%) |

**Table S11.**

Proportion of AS according to the shape of their synaptic junction within the axospinous and axodendritic synaptic population in every layer of CA1. AS: asymmetric synapses; dSP: deep part of *stratum pyramidale*; SLM: *stratum lacunosum-moleculare*; SO: *stratum oriens*; SR: *stratum radiatum*; SS: symmetric synapses; sSP: superficial part of *stratum pyramidale.*

| | Area of SAS AS ($nm^2$; mean±sem) | Perimeter of SAS AS (nm; mean±sem) | Curvature of SAS AS (mean±sem) | Area of SAS SS ($nm^2$; mean±sem) | Perimeter of SAS SS (nm; mean±sem) | Curvature of SAS SS (mean±sem) |
|---|---|---|---|---|---|---|
| SO | 86716.52±1371.02 | 1416.85±14.82 | 0.045±0.001 | 85737.60±5869.60 | 1561.04±65.55 | 0.044±0.002 |
| dSP | 92045.29±1192.92 | 1467.14±12.26 | 0.048±0.001 | 74764.69±3057.33 | 1494.82±45.84 | 0.044±0.001 |
| sSP | 88061.63±1038.49 | 1456.89±10.76 | 0.053±0.001 | 58305.43±2612.01 | 1257.36±37.28 | 0.055±0.002 |
| SR | 82841.26±1201.47 | 1389.57±12.63 | 0.052±0.001 | 63183.20±2734.96 | 1344.78±42.85 | 0.050±0.002 |
| SLM | 91419.95±1376.38 | 1477.28±14.28 | 0.044±0.001 | 57390.19±2071.04 | 1288.54±33.29 | 0.046±0.001 |
| Totals | 89727.65±5775.90 | 1458.82±56.17 | 0.050±0.002 | 67236.17±4456.52 | 1378.38±71.47 | 0.047±0.002 |

**Table S12.**

Morphological data on synapses based on synaptic apposition surface (SAS) in all layers of CA1. AS: asymmetric synapses; CA: *cornu ammonis*; dSP: deep part of *stratum pyramidale*; SD: standard deviation; sem: standard error of the mean; SLM: *stratum lacunosum-moleculare*; SO: *stratum oriens*; SR: *stratum radiatum*; SS: symmetric synapses; sSP: superficial part of *stratum pyramidale.*

| | | Area of SAS AS ($nm^2$; mean±sem) | Perimeter of SAS AS (nm; mean±sem) | Curvature of SAS AS (mean±sem) | Area of SAS SS ($nm^2$; mean±sem) | Perimeter of SAS SS (nm; mean±sem) | Curvature of SAS SS (mean±sem) |
|---|---|---|---|---|---|---|---|
| SO | Axospinous | 92900.02±4207.11 | 1490.11±42.11 | 0.045±0.001 | 76841.78±24870.78 | 1428.24±244.41 | 0.042±0.003 |
| | Axodendritic | 111843.21±15119.25 | 1626.09±127.58 | 0.045±0.004 | 88155.34±13916.54 | 1579.45±122.73 | 0.044±0.002 |
| dSP | Axospinous | 100377.53±3285.52 | 1554.92±39.94 | 0.049±0.003 | 53628.30±8432.84 | 1170.13±99.05 | 0.039±0.003 |
| | Axodendritic | 113415.07±11395.55 | 1681.01±121.94 | 0.039±0.003 | 82319.21±4124.52 | 1629.20±111.60 | 0.044±0.002 |
| sSP | Axospinous | 101533.55±10258.19 | 1588.45±93.27 | 0.053±0.001 | 45484.55±3458.52 | 995.51±56.06 | 0.048±0.007 |
| | Axodendritic | 83216.23±11310.38 | 1392.52±111.42 | 0.049±0.005 | 64386.67±9153.64 | 1327.90±108.50 | 0.053±0.003 |
| SR | Axospinous | 102259.93±11303.07 | 1586.06±101.93 | 0.054±0.001 | 52408.99±8678.45 | 1173.23±24.75 | 0.078±0.027 |
| | Axodendritic | 99896.11±23037.14 | 1514.68±205.07 | 0.047±0.003 | 60472.43±7514.73 | 1291.74±115.58 | 0.050±0.005 |
| SLM | Axospinous | 98081.18±6717.02 | 1567.25±68.94 | 0.043±0.002 | 41129.57±4778.92 | 1063.43±111.81 | 0.055±0.009 |
| | Axodendritic | 159586.41±10174.50 | 2041.64±77.33 | 0.046±0.002 | 62468.75±3482.63 | 1371.04±78.01 | 0.044±0.002 |
| Totals | Axospinous | 98200.61±6202.57 | 1548.38±63.25 | 0.050±0.001 | 49044.59±5442.64 | 1126.06±87.34 | 0.048±0.003 |
| | Axodendritic | 117360.02±8315.30 | 1686.99±82.06 | 0.045±0.002 | 71218.23±4426.35 | 1436.55±68.97 | 0.047±0.002 |

**Table S13.**

Morphological data based on synaptic apposition surface (SAS) regarding the postsynaptic targets in every layer of CA1. AS: asymmetric synapses; dSP: deep part of *stratum pyramidale*; sem: standard error of the mean; SLM: *stratum lacunosum-moleculare*; SO: *stratum oriens*; SR: *stratum radiatum*; SS: symmetric synapses; sSP: superficial part of *stratum pyramidale.*

| | | Area of AS SAS AS ($nm^2$; mean±sem) | Perimeter of SAS AS (nm; mean±sem) | Curvature of SAS AS (mean±sem) | Area of SAS SS ($nm^2$; mean±sem) | Perimeter of SAS SS (nm; mean±sem) | Curvature of SAS SS (mean±sem) |
|---|---|---|---|---|---|---|---|
| SO | Macular | 74181.28±4910.43 | 1283.10±58.28 | 0.043±0.002 | 72168.70±10458.36 | 1330.32±104.11 | 0.043±0.002 |
| | Horseshoe-shaped | 171171.18±8988.19 | 2787.25±140.87 | 0.065±0.006 | 101225.81±10559.25 | 2078.29±213.47 | 0.041±0.004 |
| | Perforated | 188978.45±13785.62 | 2426.28±124.62 | 0.055±0.008 | 233367.89±77405.62 | 3306.30±766.80 | 0.064±0.022 |
| | Fragmented | 223086.00±10491.93 | 2542.75±180.04 | 0.099±0.015 | 167220.54±41188.75 | 2187.00±204.15 | 0.064±0.006 |
| dSP | Macular | 73347.29±1957.92 | 1272.02±18.33 | 0.044±0.002 | 58981.36±3424.93 | 1244.50±56.04 | 0.042±0.002 |
| | Horseshoe-shaped | 185893.89±12000.00 | 3046.28±122.58 | 0.086±0.008 | 114528.91±12807.42 | 2300.59±160.87 | 0.044±0.005 |
| | Perforated | 205732.10±9992.43 | 2560.79±90.50 | 0.062±0.003 | 114415.03±16821.37 | 2071.89±204.10 | 0.047±0.005 |
| | Fragmented | 274778.26±18518.33 | 2368.79±142.38 | 0.120±0.013 | 76662.93±11428.32 | 1277.38±191.22 | 0.065±0.022 |
| sSP | Macular | 70946.54±5240.96 | 1276.40±50.50 | 0.048±0.001 | 51365.93±5150.52 | 1132.33±53.80 | 0.053±0.002 |
| | Horseshoe-shaped | 213184.36±24896.05 | 3229.59±262.31 | 0.092±0.008 | 99774.37±13522.27 | 2135.95±172.18 | 0.046±0.011 |
| | Perforated | 225076.44±26339.64 | 2759.96±232.75 | 0.078±0.003 | 118372.40±38494.85 | 2139.19±366.81 | 0.031±0.005 |
| | Fragmented | 266724.88±29303.55 | 2563.51±172.67 | 0.131±0.017 | 110887.92±32290.73 | 1600.21±309.21 | 0.071±0.003 |
| SR | Macular | 65852.36±3067.58 | 1205.12±25.97 | 0.048±0.004 | 51088.37±4910.94 | 1128.12±54.26 | 0.051±0.006 |
| | Horseshoe-shaped | 181125.70±17161.52 | 2854.53±230.13 | 0.071±0.003 | 105630.16±11089.39 | 2147.72±257.86 | 0.034±0.010 |
| | Perforated | 201824.60±23993.63 | 2638.11±248.50 | 0.076±0.004 | 63548.92±27451.37 | 1328.90±466.08 | 0.044±0.013 |
| | Fragmented | 269475.34±35799.23 | 2683.22±317.04 | 0.118±0.014 | 89588.42±11515.62 | 1563.85±63.03 | 0.049±0.003 |
| SLM | Macular | 72110.84±2661.42 | 1247.41±16.51 | 0.041±0.002 | 50704.33±1595.42 | 1184.14±33.54 | 0.047±0.003 |
| | Horseshoe-shaped | 160347.91±12123.34 | 2673.00±86.45 | 0.054±0.003 | 65798.60±10255.21 | 1630.59±168.00 | 0.041±0.003 |
| | Perforated | 201177.54±15526.48 | 2542.94±79.43 | 0.057±0.002 | 95746.01±14562.01 | 1933.97±218.85 | 0.036±0.004 |
| | Fragmented | 202913.98±8401.89 | 2199.00±87.97 | 0.077±0.009 | 149534.82±11567.29 | 1834.22±244.95 | 0.062±0.002 |

| | | | | | | | |
|---|---|---|---|---|---|---|---|
| Totals | Macular | 70322.92±2524.70 | 1251.59±27.36 | 0.045±0.002 | 56769.81±2226.39 | 1204.01±37.92 | 0.047±0.003 |
| | Horseshoe-shaped | 146474.80±8979.65 | 2908.08±132.62 | 0.075±0.003 | 100178.06±4541.53 | 2132.38±75.35 | 0.044±0.003 |
| | Perforated | 205500.73±13875.57 | 2598.53±137.87 | 0.067±0.001 | 135419.88±24500.18 | 2206.83±210.21 | 0.047±0.005 |
| | Fragmented | 249642.43±17666.73 | 2480.97±157.13 | 0.091±0.021 | 109912.28±12528.19 | 1655.41±92.67 | 0.062±0.003 |

**Table S14.**

Morphological data based on synaptic apposition surface (SAS) of synapses regarding the shape of the synaptic junction in every layer of CA1. AS: asymmetric synapses; dSP: deep part of *stratum pyramidale*; SAS: synaptic apposition surface; sem: standard error of the mean; SLM: *stratum lacunosum-moleculare*; SO: *stratum oriens*; SR: *stratum radiatum*; SS: symmetric synapses; sSP: superficial part of *stratum pyramidale.*

| | | Thickness (mm; mean±SD) | Percentage of all CA1 layers | $V_{bv}$ (%; mean±SD) | $V_g$ (%; mean±SD) | $V_{neu}$ (%; mean±SD) | $V_n$ (%; mean±SD) |
|---|---|---|---|---|---|---|---|
| AB1 | SO | 0.07±0.02 | 2% | 13.03±0.59 | 1.54±0.70 | 0.14±0.24 | 85.30±1.26 |
| | SP | 1.66±0.03 | 49% | 6.10±0.33 | 1.03±0.26 | 3.80±0.35 | 89.07±0.20 |
| | SR | 0.47±0.11 | 14% | 4.57±0.48 | 2.19±0.90 | 0.04±0.08 | 93.20±1.15 |
| | SLM | 0.79±0.04 | 23% | 5.38±0.91 | 2.03±0.51 | 0.03±0.04 | 92.57±1.40 |
| | All layers | 3.41±0.06 | 100% | - | - | - | - |
| AB2 | SO | 0.03±0.01 | 1% | 8.15±2.99 | 2.32±0.71 | 0.00±0.00 | 89.53±2.52 |
| | SP | 1.07±0.03 | 47% | 4.88±0.33 | 0.60±0.31 | 4.94±0.39 | 89.57±0.31 |
| | SR | 0.27±0.01 | 12% | 6.04±1.21 | 0.80±0.52 | 0.44±0.22 | 92.73±1.42 |
| | SLM | 0.62±0.05 | 27% | 7.10±0.36 | 0.36±0.31 | 0.03±0.05 | 92.51±0.68 |
| | All layers | 2.27±0.03 | 100% | - | - | - | - |
| AB3 | SO | 0.04±0.01 | 2% | 5.50±1.35 | 3.24±1.87 | 1.16±1.09 | 90.10±1.85 |
| | SP | 0.82±0.11 | 39% | 4.02±0.81 | 0.73±0.23 | 5.69±1.50 | 89.56±2.44 |
| | SR | 0.40±0.06 | 19% | 4.20±0.96 | 0.56±0.50 | 0.28±0.09 | 94.97±0.45 |
| | SLM | 0.43±0.05 | 20% | 4.46±1.37 | 1.67±0.40 | 0.00±0.00 | 93.87±1.08 |
| | All layers | 2.13±0.08 | 100% | - | - | - | - |
| AB4 | SO | 0.10±0.01 | 4% | 6.42±2.74 | 1.38±0.28 | 0.69±0.72 | 91.51±2.99 |
| | SP | 0.90±0.19 | 38% | 6.08±0.19 | 0.26±0.17 | 3.71±0.58 | 89.95±0.73 |
| | SR | 0.55±0.08 | 24% | 4.89±0.93 | 0.53±0.46 | 0.00±0.00 | 94.58±1.34 |
| | SLM | 0.62±0.01 | 26% | 7.40±1.13 | 0.29±0.06 | 0.08±0.13 | 92.23±1.20 |
| | All layers | 2.34±0.08 | 100% | - | - | - | - |
| M17 | SO | 0.05±0.00 | 1% | 4.81±0.59 | 1.27±0.17 | 0.30±0.07 | 93.63±0.61 |
| | SP | 1.17±0.11 | 35% | 4.21±1.69 | 0.37±0.17 | 3.01±0.30 | 92.41±1.85 |
| | SR | 1.07±0.18 | 32% | 4.24±0.48 | 0.31±0.27 | 0.00±0.00 | 95.46±0.68 |
| | SLM | 0.67±0.03 | 20% | 5.47±1.71 | 0.88±0.19 | 0.05±0.09 | 93.60±1.83 |
| | All layers | 3.35±0.23 | 100% | - | - | - | - |

**Table S15.**

Data on CA1 thickness and volume fraction occupied by different cortical elements per case. CA: *cornu ammonis*; SD: standard deviation; SLM: *stratum lacunosum-moleculare*; SO: *stratum oriens*; SP: *stratum pyramidale*; SR: *stratum radiatum*; $V_{bv}$: volume fraction occupied by blood vessels; $V_g$: volume fraction occupied by glia; $V_{neu}$: volume fraction occupied by neurons; $V_n$: volume fraction occupied by neuropil.

The following significant differences were observed between cases:

- SO:
    - $V_{bv}$:
        - AB1-AB3 (ANOVA, p=0.005)
        - AB1-AB4 (ANOVA, p=0.013)
        - AB1-M17 (ANOVA, p=0.003)
    - $V_n$:
        - AB1-AB4 (ANOVA, p=0.024)
        - AB1-M17 (ANOVA, p=0.004)
- SP:
    - $V_{neu}$:
        - AB3-M17 (ANOVA, p=0.011)
- SR:
    - $V_{neu}$:
        - AB1-AB2 (ANOVA, p=0.004)
        - AB2-AB4 (ANOVA, p=0.002)
        - AB2-M17 (ANOVA, p=0.002)
        - AB3-AB4 (ANOVA, p=0.039)
        - AB3-M17 (ANOVA, p=0.039)
    - $V_{glia}$:
        - AB1-AB2 (ANOVA, p=0.082)
        - AB1-AB3 (ANOVA, p=0.038)
        - AB1-AB4 (ANOVA, p=0.034)
        - AB1-M17 (ANOVA, p=0.015)
- SLM:
    - $V_{glia}$:
        - AB1-AB2 (ANOVA, p=0.001)
        - AB1-AB4 (ANOVA, p=0.001)
        - AB1-M17 (ANOVA, p=0.019)
        - AB2-AB3 (ANOVA, p=0.006)
        - AB3-AB4 (ANOVA, p=0.004).

| *STRATUM ORIENS* | | | | | |
|---|---|---|---|---|---|
| | AB1 | AB2 | AB3 | AB4 | M17 |
| No. AS | 399 | 961 | 446 | 346 | 496 |
| No. SS | 20 | 48 | 40 | 15 | 43 |
| % AS | 95.23% | 95.24% | 91.77% | 95.84% | 92.02% |
| % SS | 4.77% | 4.76% | 8.23% | 4.16% | 7.98% |
| CF volume ($\mu m^3$) | 1085 | 1281 | 1126 | 1308 | 1422 |
| Density AS/$\mu m^3$ (mean±SD) | 0.37±0.04 | 0.75±0.16 | 0.40±0.11 | 0.27±0.06 | 0.35±0.05 |
| Density SS/$\mu m^3$ (mean±SD) | 0.02±0.00 | 0.04±0.01 | 0.03±0.02 | 0.01±0.01 | 0.03±0.01 |
| Density AS+SS/$\mu m^3$ (mean±SD) | 0.39± 0.04 | 0.78± 0.17 | 0.43± 0.10 | 0.28± 0.07 | 0.38± 0.04 |
| SAS Area of AS ($nm^2$; mean±sem) | 93342.82± 3268.05 | 73718.73± 1964.31 | 94969.69± 3940.23 | 107520.65± 4444.53 | 84635.58± 2937.08 |
| SAS Perimeter of AS (nm; mean±sem) | 1438.33± 36.58 | 1267.48± 21.07 | 1513.43± 44.28 | 1619.57± 42.70 | 1460.72± 33.10 |
| SAS Curvature of AS (mean±sem) | 0.046± 0.001 | 0.044± 0.001 | 0.046± 0.002 | 0.051± 0.002 | 0.040± 0.001 |
| SAS Area of SS ($nm^2$; mean±sem) | 70229.23± 10794.99 | 61411.26± 6274.37 | 139459.24± 18604.16 | 88163.18± 18312.31 | 68925.38± 4248.17 |
| SAS Perimeter of SS (nm; mean±sem) | 1341.18± 142.26 | 1332.72± 96.05 | 2039.02± 183.62 | 1552.94± 235.24 | 1471.26± 79.48 |
| SAS Curvature of SS (mean±sem) | 0.048± 0.004 | 0.045± 0.003 | 0.047± 0.007 | 0.045± 0.006 | 0.037± 0.002 |
| Distance to nearest synapse (nm; mean±SD) | 798.45± 71.69 | 647.61± 48.08 | 732.96± 48.85 | 802.36± 37.28 | 732.66± 11.24 |

**Table S16.**
Ultrastructural analysis of the neuropil of the *stratum oriens* of CA1 per case. AS: asymmetric synapses; CF: counting frame; No.: number; SAS: synaptic apposition surface; SD: standard deviation; sem: standard error of the mean; SS: symmetric synapses.

The following significant differences were observed between cases:

- Synaptic density:
    - AB1-AB2 (ANOVA, p=0.004)
    - AB2-AB3 (ANOVA, p=0.008)
    - AB2-AB4 (ANOVA, p=0.003)
    - AB2-M17 (ANOVA, p=0.001)
- AS SAS Area:
    - AB2-AB4 (ANOVA, p=0.004)
    - AB4-M17 (ANOVA, p=0.031)
- AS SAS Perimeter:
    - AB2-AB4 (ANOVA, p=0.016)
- AS SAS Curvature:
    - AB4-M17 (ANOVA, p=0.009).

| *STRATUM PYRAMIDALE* (DEEP) | | | | | |
|---|---|---|---|---|---|
| | AB1 | AB2 | AB3 | AB4 | M17 |
| No. AS | 528 | 1168 | 694 | 708 | 751 |
| No. SS | 45 | 103 | 39 | 46 | 48 |
| % AS | 92.15% | 91.90% | 94.68% | 93.90% | 93.99% |
| % SS | 7.85% | 8.10% | 5.32% | 6.10% | 6.01% |
| CF volume ($\mu m^3$) | 1181 | 1241 | 1061 | 1281 | 1239 |
| Density of AS/$\mu m^3$ (mean±SD) | 0.48±0.06 | 1.05±0.41 | 0.69±0.01 | 0.58±0.05 | 0.65±0.26 |
| Density of SS/$\mu m^3$ (mean±SD) | 0.04±0.01 | 0.08±0.03 | 0.04±0.01 | 0.04±0.00 | 0.04±0.02 |
| Density of AS+SS/$\mu m^3$ (mean±SD) | 0.48± 0.06 | 1.05± 0.41 | 0.69± 0.01 | 0.58± 0.05 | 0.65± 0.26 |
| SAS Area of AS ($nm^2$; mean±sem) | 92722.75± 2681.12 | 81061.60± 2024.49 | 97161.24± 3191.30 | 101859.21± 2938.24 | 94678.63± 2678.52 |
| SAS Perimeter of AS (nm; mean±sem) | 1453.48± 27.87 | 1333.65± 19.93 | 1504.59± 29.95 | 1605.18± 32.69 | 1519.67± 28.75 |
| SAS Curvature of AS (mean±sem) | 0.045± 0.001 | 0.043± 0.001 | 0.056± 0.002 | 0.057± 0.001 | 0.044± 0.001 |
| SAS Area of SS ($nm^2$; mean±sem) | 67643.28± 7427.96 | 72582.05± 4969.33 | 91389.02± 10215.35 | 69230.11± 5762.38 | 77574.95± 7439.81 |
| SAS Perimeter of SS (nm; mean±sem) | 1236.47± 94.11 | 1433.30± 65.90 | 1870.83± 161.56 | 1428.71± 86.55 | 1619.03± 128.33 |
| SAS Curvature of SS (mean±sem) | 0.041± 0.004 | 0.047± 0.002 | 0.044± 0.003 | 0.046± 0.003 | 0.037± 0.002 |
| Distance to nearest synapse (nm; mean±SD) | 722.14± 27.54 | 586.62± 59.88 | 646.90± 5.95 | 704.29± 4.97 | 689.09± 93.50 |

**Table S17.**
Ultrastructural analysis of the neuropil of the deep part of *stratum pyramidale* of CA1 per case. AS: asymmetric synapses; CF: counting frame; SAS: synaptic apposition surface; SD: standard deviation; sem: standard error of the mean; SS: symmetric synapses.

The following significant differences were observed between cases:

- AS SAS Perimeter:
  - AB2-AB4 (ANOVA, p=0.017)
- AS SAS Curvature:
  - AB2-AB4 (ANOVA, p=0.043)
  - AB4-M17 (ANOVA, p=0.043).

| *STRATUM PYRAMIDALE* (SUP) | | | | | |
|---|---|---|---|---|---|
| | AB1 | AB2 | AB3 | AB4 | M17 |
| No. AS | 1063 | 1242 | 1180 | 697 | 1001 |
| No. SS | 31 | 43 | 30 | 49 | 43 |
| % AS | 97.17% | 96.65% | 97.52% | 93.43% | 95.88% |
| % SS | 2.83% | 3.35% | 2.48% | 6.57% | 4.12% |
| CF volume (μm$^3$) | 1008 | 1125 | 1114 | 1054 | 1100 |
| Density of AS/μm$^3$ (mean±SD) | 1.06±.220.0 8 | 1.11±0.22 | 1.06±0.14 | 0.66±0.10 | 0.90±0.07 |
| Density of SS/μm$^3$ (mean±SD) | 0.03±0.01 | 0.04±0.01 | 0.03±0.01 | 0.05±0.01 | 0.04±0.01 |
| Density of AS+SS/μm$^3$ (mean±SD) | 1.09± 0.08 | 1.14± 0.23 | 1.09± 0.13 | 0.71± 0.10 | 0.94± 0.08 |
| SAS Area of AS (nm$^2$; mean±sem) | 70955.73± 1513.22 | 78494.59± 1949.28 | 81458.42± 1967.25 | 120382.77± 3462.29 | 103428.73± 2809.43 |
| SAS Perimeter of AS (nm; mean±sem) | 1327.93± 17.92 | 1325.21± 19.52 | 1405.83± 20.29 | 1726.02± 34.42 | 1630.59± 29.97 |
| SAS Curvature of AS (mean±sem) | 0.054± 0.001 | 0.054± 0.001 | 0.053± 0.001 | 0.058± 0.001 | 0.050± 0.001 |
| SAS Area of SS (nm$^2$; mean±sem) | 42279.64± 4088.94 | 44082.07± 4942.68 | 73446.84± 12905.52 | 76579.79± 10317.67 | 64714.08± 7793.82 |
| SAS Perimeter of SS (nm; mean±sem) | 1072.17± 61.40 | 1110.70± 93.53 | 1511.62± 182.48 | 1387.66± 126.13 | 1283.53± 94.11 |
| SAS Curvature of SS (mean±sem) | 0.048± 0.004 | 0.059± 0.005 | 0.060± 0.005 | 0.052± 0.003 | 0.046± 0.004 |
| Distance to nearest synapse (nm; mean±SD) | 589.48± 37.31 | 571.68± 25.00 | 579.05± 11.49 | 665.82± 53.95 | 613.96± 28.69 |

**Table S18.**

Ultrastructural analysis of the neuropil of the superficial part of *stratum pyramidale* of CA1 per case. AS: asymmetric synapses; CF: counting frame; SAS: synaptic apposition surface; SD: standard deviation; sem: standard error of the mean; SS: symmetric synapses; Sup: superficial.

The following significant differences were observed between cases:

- AS:SS ratio:
  - AB1-AB4 ($\chi^2$, p=0.0002)
  - AB3-AB4 ($\chi^2$, p=$1.475 \times 10^{-5}$)
- AS SAS Area:
  - AB1-AB4 (ANOVA, p=0.0002)
  - AB1-M17 (ANOVA, p=0.006)
  - AB2-AB4 (ANOVA, p=0.0009)
  - AB2-M17 (ANOVA, p=0.036)
  - AB3-AB4 (ANOVA, p=0.002)
- AS SAS Perimeter:
  - AB1-AB4 (ANOVA, p=$6.182 \times 10^{-5}$)
  - AB1-M17 (ANOVA, p=0.0005)
  - AB2-AB4 (ANOVA, p=$6.351 \times 10^{-5}$)
  - AB2-M17 (ANOVA, p=0.0006)
  - AB3-AB4 (ANOVA, p=0.0004)
  - AB3-M17 (ANOVA, p=0.006).

| *STRATUM RADIATUM* | | | | | |
|---|---|---|---|---|---|
| | AB1 | AB2 | AB3 | AB4 | M17 |
| No. AS | 647 | 897 | 1048 | 572 | 672 |
| No. SS | 25 | 24 | 39 | 42 | 42 |
| % AS | 96.28% | 97.39% | 96.41% | 93.16% | 94.12% |
| % SS | 3.72% | 2.61% | 3.59% | 6.84% | 5.88% |
| CF volume (µm$^3$) | 1310 | 1075 | 1176 | 1085 | 1362 |
| Density of AS/µm$^3$ (mean±SD) | 0.50±0.03 | 0.83±0.10 | 0.87±0.17 | 0.53±0.11 | 0.49±0.04 |
| Density of SS/µm$^3$ (mean±SD) | 0.02±0.01 | 0.02±0.02 | 0.03±0.02 | 0.04±0.01 | 0.03±0.01 |
| Density of AS+SS/µm$^3$ (mean±SD) | 0.52± 0.03 | 0.85± 0.09 | 0.91± 0.16 | 0.57± 0.12 | 0.53± 0.05 |
| SAS Area of AS (nm$^2$; mean±sem) | 76444.31± 2197.02 | 72546.21± 2322.56 | 72385.57± 1854.08 | 105184.59± 3515.39 | 100176.82± 1201.47 |
| SAS Perimeter of AS (nm; mean±sem) | 1338.18± 22.34 | 1295.34± 25.94 | 1299.66± 20.15 | 1581.08± 36.93 | 1543.33± 38.14 |
| SAS Curvature of AS (mean±sem) | 0.052± 0.001 | 0.056± 0.001 | 0.048± 0.001 | 0.062± 0.002 | 0.043± 0.001 |
| SAS Area of SS (nm$^2$; mean±sem) | 52100.88± 8050.33 | 41303.20± 6506.56 | 53992.48± 7792.31 | 64738.80± 6049.48 | 86398.68± 7593.45 |
| SAS Perimeter of SS (nm; mean±sem) | 1156.15± 115.84 | 1010.78± 107.58 | 1177.10± 109.14 | 1361.28± 116.76 | 1677.58± 125.05 |
| SAS Curvature of SS (mean±sem) | 0.045± 0.004 | 0.065± 0.009 | 0.050± 0.005 | 0.057± 0.005 | 0.035± 0.003 |
| Distance to nearest synapse (nm; mean±SD) | 694.96± 15.37 | 600.94± 8.77 | 587.65± 46.61 | 724.52± 31.04 | 701.98± 43.51 |

**Table S19.**
Ultrastructural analysis of the neuropil of the *stratum radiatum* of CA1 per case. AS: asymmetric synapses; CF: counting frame; SAS: synaptic apposition surface; SD: standard deviation; sem: standard error of the mean; SS: symmetric synapses.

The following significant differences were observed between cases:

- Synaptic density:
  - AB1-AB2 (ANOVA, p=0.017)
  - AB1-AB3 (ANOVA, p=0.007)
  - AB2-AB4 (ANOVA, p=0.021)
  - AB2-M17 (ANOVA, p=0.044)
  - AB3-AB4 (ANOVA, p=0.008)
  - AB3-M17 (ANOVA, p=0.017)
- AS:SS ratio:
  - AB2-AB4 ($\chi^2$, p=9.165x10$^{-5}$)
  - AB2-M17 ($\chi^2$, p=0.0009)
- AS SAS Area:
  - AB1-AB4 (ANOVA, p=0.012)
  - AB1-M17 (ANOVA, p=0.045)
  - AB2-AB4 (ANOVA, p=0.004)
  - AB2-M17 (ANOVA, p=0.015)
  - AB3-AB4 (ANOVA, p=0.004)
  - AB3-M17 (ANOVA, p=0.015)
- AS SAS Perimeter:
  - AB1-AB4 (ANOVA, p=0.039)
  - AB2-AB4 (ANOVA, p=0.014)
  - AB2-M17 (ANOVA, p=0.038)
  - AB3-AB4 (ANOVA, p=0.016)
  - AB3-M17 (ANOVA, p=0.046)
- AS SAS Curvature:
  - AB1-AB4 (ANOVA, p=0.033)
  - AB2-M17 (ANOVA, p=0.008)
  - AB3-AB4 (ANOVA, p=0.006)
  - AB4-M17 (ANOVA, p=0.0005)
- SS SAS Area:
  - AB2-AB4 (ANOVA, p=0.030)
  - AB2-M17 (ANOVA, p=0.010).

| *STRATUM LACUNOSUM-MOLECULARE* | | | | | |
|---|---|---|---|---|---|
| | AB1 | AB2 | AB3 | AB4 | M17 |
| No. AS | 455 | 533 | 702 | 429 | 503 |
| No. SS | 43 | 40 | 70 | 70 | 93 |
| % AS | 91.37% | 93.02% | 90.93% | 85.97% | 84.40% |
| % SS | 8.63% | 6.98% | 9.07% | 14.03% | 15.60% |
| CF volume (μm$^3$) | 1141 | 1127 | 1278 | 1035 | 1109 |
| Density of AS/μm$^3$ (mean±SD) | 0.40±0.05 | 0.47±0.04 | 0.56±0.15 | 0.39±0.07 | 0.49±0.04 |
| Density of SS/μm$^3$ (mean±SD) | 0.04±0.00 | 0.04±0.01 | 0.05±0.01 | 0.06±0.01 | 0.09±0.01 |
| Density of AS+SS/μm$^3$ (mean±SD) | 0.44± 0.05 | 0.51± 0.04 | 0.61± 0.15 | 0.45± 0.07 | 0.58± 0.04 |
| SAS Area of AS (nm$^2$; mean±sem) | 88303.06± 3115.26 | 98697.11± 3522.76 | 76680.19± 2143.59 | 112152.43± 4038.28 | 89394.31± 1376.38 |
| SAS Perimeter of AS (nm; mean±sem) | 1149.67± 33.63 | 1507.70± 32.98 | 1375.86± 23.90 | 1604.79± 39.69 | 1502.69± 32.95 |
| SAS Curvature of AS (mean±sem) | 0.047± 0.002 | 0.042± 0.001 | 0.047± 0.001 | 0.044± 0.002 | 0.040± 0.001 |
| SAS Area of SS (nm$^2$; mean±sem) | 47017.73± 4264.25 | 72916.74± 7087.22 | 61925.95± 5471.59 | 53253.73± 2769.68 | 59848.75± 3890.73 |
| SAS Perimeter of SS (nm; mean±sem) | 1086.19± 63.88 | 1589.32± 129.80 | 1446.40± 93.66 | 1191.99± 43.82 | 1287.05± 53.39 |
| SAS Curvature of SS (mean±sem) | 0.046± 0.003 | 0.045± 0.003 | 0.050± 0.002 | 0.053± 0.004 | 0.038± 0.002 |
| Distance to nearest synapse (nm; mean±SD) | 696.49± 22.38 | 715.96± 52.01 | 651.23± 49.79 | 695.65± 20.83 | 688.92± 31.36 |

**Table S20.**
Ultrastructural analysis of the neuropil of the *stratum lacunosum-moleculare* of CA1 per case. AS: asymmetric synapses; CF: counting frame; SAS: synaptic apposition surface; SD: standard deviation; sem: standard error of the mean; SS: symmetric synapses.

The following significant differences were observed between cases:

- AS:SS ratio:
    - AB1-M17 ($\chi^2$, p=0.0005)
    - AB2-AB4 ($\chi^2$, p=0.0002)
    - AB2-M17 ($\chi^2$, p=3.332x$10^{-6}$)
    - AB3-M17($\chi^2$, p=0.0003)
- AS SAS Area:
    - AB3-AB4 (ANOVA, p=0.006)
- SS SAS Area:
    - AB1-AB2 (ANOVA, p=0.026)
- SS SAS Perimeter:
    - AB1-AB2 (ANOVA, p=0.002)
    - AB1-AB3 (ANOVA, p=0.015)
    - AB2-AB4 (ANOVA, p=0.009)
    - AB2-M17 (ANOVA, p=0.048).

| *STRATUM ORIENS* | | | | | | |
|---|---|---|---|---|---|---|
| | | No. AS | No. SS | SAS Area of AS ($nm^2$; mean±sem) | SAS Perimeter of AS (nm; mean±sem) | SAS Curvature of AS (mean±sem) |
| Axospinous | AB1 | 254 (77.44%) | 1 (0.30%) | 97732.76±4179.97 | 1484.16±46.83 | 0.045±0.002 |
| | AB2 | 455 (87.33%) | 4 (0.77%) | 81382.46±3033.28 | 1349.53±32.93 | 0.043±0.001 |
| | AB3 | 226 (73.14%) | 2 (0.65%) | 94628.46±5446.79 | 1535.91±65.99 | 0.045±0.002 |
| | AB4 | 184 (83.26%) | 2 (0.90%) | 104936.32±5177.26 | 1605.67±56.16 | 0.048±0.002 |
| | M17 | 240 (60.30%) | 0 (0.00%) | 85820.10±4478.71 | 1475.30±52.95 | 0.042±0.002 |
| Axodendritic | AB1 | 56 (17.07%) | 17 (5.18%) | 98681.86±7172.21 | 1449.85±66.75 | 0.040±0.004 |
| | AB2 | 22 (4.22%) | 40 (7.68%) | 69026.21±11540.48 | 1233.45±145.04 | 0.048±0.006 |
| | AB3 | 43 (13.92%) | 38 (12.30%) | 128191.12±13286.59 | 1806.62±135.43 | 0.044±0.003 |
| | AB4 | 22 (9.95%) | 13 (5.88%) | 159160.95±32555.49 | 1947.36±191.55 | 0.059±0.009 |
| | M17 | 116 (29.15%) | 42 (10.55%) | 104155.92±5740.68 | 1693.18±60.24 | 0.035±0.002 |

**Table S21.**
Ultrastructural data regarding the postsynaptic target in the *stratum oriens* of CA1 per case. AS: asymmetric synapses; SAS: synaptic apposition surface; sem: standard error of the mean; SS: symmetric synapses.

The following significant differences were observed between cases:

- AB1-AB2 ($\chi^2$, p=6.119x$10^{-9}$)
  - Axospinous AS ($\chi^2$, p=0.0002)
  - Axodendritic AS ($\chi^2$, p=5.906x$10^{-10}$)
- AB1-M17 ($\chi^2$, p=7.802x$10^{-6}$)
  - Axospinous AS ($\chi^2$, p=8.300x$10^{-7}$)
  - Axodendritic AS ($\chi^2$, p=0.0002)

- AB2-AB3 ($\chi^2$, p=4.150x10$^{-7}$)
    - Axospinous AS ($\chi^2$, p=5.445x10$^{-7}$)
    - Axodendritic AS ($\chi^2$, p=1.509x10$^{-6}$)

- AB2-M17 ($\chi^2$, p=1.000x10$^{-17}$)
    - Axospinous AS ($\chi^2$, p=1.000x10$^{-17}$)
    - Axodendritic AS ($\chi^2$, p=1.000x10$^{-17}$)
- AB3-M17 ($\chi^2$, p=1.315x10$^{-5}$)
    - Axospinous AS ($\chi^2$, p=0.0004)
    - Axodendritic AS ($\chi^2$, p=1.274x10$^{-6}$)
- AB4-M17 ($\chi^2$, p=5.130x10$^{-9}$)
    - Axospinous AS ($\chi^2$, p=1.832x10$^{-9}$)
    - Axodendritic AS ($\chi^2$, p=1.155x10$^{-8}$).

| *STRATUM PYRAMIDALE* (DEEP) | | | | | | |
|---|---|---|---|---|---|---|
| | | No. AS | No. SS | SAS Area of AS ($nm^2$; mean±sem) | SAS Perimeter of AS (nm; mean±sem) | SAS Curvature of AS (mean±sem) |
| Axospinous | AB1 | 417 (86.69%) | 14 (2.91%) | 95332.59±2932.72 | 1479.39±30.01 | 0.045±0.001 |
| | AB2 | 357 (67.61%) | 6 (1.14%) | 92357.50±3763.36 | 1452.66±37.06 | 0.044±0.001 |
| | AB3 | 347 (85.68%) | 4 (0.99%) | 107947.49±4698.89 | 1609.92±45.61 | 0.056±0.002 |
| | AB4 | 334 (84.56%) | 3 (0.76%) | 97942.40±3753.44 | 1565.42±41.13 | 0.057±0.002 |
| | M17 | 333 (84.52%) | 10 (2.54%) | 108307.66±4306.91 | 1667.19±47.13 | 0.045±0.002 |
| Axodendritic | AB1 | 23 (4.78%) | 27 (5.61%) | 80973.53±14337.30 | 1356.75±147.81 | 0.037±0.004 |
| | AB2 | 121 (22.92%) | 44 (8.33%) | 108858.10±7942.19 | 1563.64±64.53 | 0.043±0.002 |
| | AB3 | 22 (5.43%) | 32 (7.90%) | 107971.89±17989.34 | 1591.62±172.83 | 0.038±0.005 |
| | AB4 | 20 (5.06%) | 38 (9.62%) | 151864.86±20111.74 | 2067.97±192.79 | 0.047±0.006 |
| | M17 | 16 (4.06%) | 35 (8.88%) | 117406.98±20556.63 | 1825.06±192.09 | 0.031±0.003 |

**Table S22.**
Ultrastructural data regarding the postsynaptic target in the deep part of the *stratum pyramidale* of CA1 per case. AS: asymmetric synapses; SAS: synaptic apposition surface; sem: standard error of the mean; SS: symmetric synapses.

The following significant differences were observed between cases:

- AB1-AB2 ($\chi^2$, p=1.00x$10^{-17}$)
  - Axospinous AS ($\chi^2$, 4.666x$10^{-13}$)
  - Axodendritic AS ($\chi^2$, p=1.00x$10^{-17}$)
- AB2-AB3 ($\chi^2$, p=4.801x$10^{-12}$)
  - Axospinous AS ($\chi^2$, p=1.301x$10^{-10}$)
  - Axodendritic AS ($\chi^2$, p=2.400x$10^{-15}$)

- AB2-AB4 ($\chi^2$, $p=3.187\times10^{-12}$)
    - Axospinous AS ($\chi^2$, $p=2.804\times10^{-9}$)
    - Axodendritic AS ($\chi^2$, $p=6.000\times10^{-16}$)

- AB2-M17 ($\chi^2$, $p=4.500\times10^{-15}$)
    - Axospinous AS ($\chi^2$, $p=2.899\times10^{-9}$)
    - Axodendritic AS ($\chi^2$, $p=1.000\times10^{-17}$).

| *STRATUM PYRAMIDALE* (SUP) | | | | | | |
|---|---|---|---|---|---|---|
| | | No. AS | No. SS | SAS Area of AS ($nm^2$; mean±sem) | SAS Perimeter of AS (nm; mean±sem) | SAS Curvature of AS (mean±sem) |
| Axospinous | AB1 | 772 (92.12%) | 3 (0.36%) | 73989.28±1751.81 | 1358.47±20.92 | 0.054±0.001 |
| | AB2 | 314 (83.96%) | 0 (0.00%) | 88278.10±4154.22 | 1426.42±42.29 | 0.050±0.002 |
| | AB3 | 389 (84.93%) | 0 (0.00%) | 95024.61±3806.30 | 1548.99±39.25 | 0.053±0.002 |
| | AB4 | 314 (82.41%) | 6 (1.57%) | 127284.99±5364.09 | 1794.53±53.55 | 0.057±0.002 |
| | M17 | 395 (89.37%) | 8 (1.81%) | 123090.76±5013.11 | 1813.85±52.07 | 0.052±0.002 |
| Axodendritic | AB1 | 36 (4.30%) | 27 (3.22%) | 67033.19±8260.31 | 1225.02±79.48 | 0.052±0.005 |
| | AB2 | 26 (6.95%) | 34 (9.09%) | 49507.36±8780.47 | 1062.92±131.20 | 0.057±0.007 |
| | AB3 | 41 (8.95%) | 28 (6.11%) | 95079.45±12194.84 | 1548.29±112.02 | 0.049±0.005 |
| | AB4 | 21 (5.51%) | 40 (10.50%) | 114406.11±14319.52 | 1683.78±154.54 | 0.054±0.009 |
| | M17 | 7 (1.58%) | 32 (7.24%) | 90055.03±31084.26 | 1442.59±225.25 | 0.031±0.001 |

**Table S23.**
Ultrastructural data regarding the postsynaptic target in the superficial part of the *stratum pyramidale* of CA1 per case. AS: asymmetric synapses; SAS: synaptic apposition surface; sem: standard error of the mean; SS: symmetric synapses.

The following significant differences were observed between cases:

- AB1-AB2 ($\chi^2$, p=1.939x$10^{-5}$)
  - Axospinous AS ($\chi^2$, p=3.815x$10^{-5}$)
- AB1-AB3 ($\chi^2$, p= 0.0003)
  - Axospinous AS ($\chi^2$, p=0.0001)
- AB1-AB4 ($\chi^2$, p=1.901x$10^{-7}$)
  - Axospinous AS ($\chi^2$, p=1.414x$10^{-6}$)

- AB2-M17 ($\chi^2$, $p=4.564x10^{-5}$)
    - Axodendritic AS ($\chi^2$, $p=0.0001$)
- AB-M17 ($\chi^2$, $p=7.562x10^{-7}$)
    - Axospinous AS ($\chi^2$, $p=6.692x10^{-7}$).

| *STRATUM RADIATUM* | | | | | | |
|---|---|---|---|---|---|---|
| | | No. AS | No. SS | SAS Area of AS ($nm^2$; mean±sem) | SAS Perimeter of AS (nm; mean±sem) | SAS Curvature of AS (mean±sem) |
| Axospinous | AB1 | 406 (87.31%) | 4 (0.86%) | 78615.15±2747.63 | 1369.75±28.28 | 0.052±0.002 |
| | AB2 | 159 (67.09%) | 1 (0.42%) | 83659.03±7027.88 | 1414.65±75.42 | 0.056±0.003 |
| | AB3 | 326 (80.49%) | 0 (0.00%) | 91889.20±4501.61 | 1513.55±48.31 | 0.054±0.002 |
| | AB4 | 188 (72.31%) | 0 (0.00%) | 119670.82±6464.90 | 1714.99±67.55 | 0.058±0.002 |
| | M17 | 199 (75.38%) | 1 (0.38%) | 137465.45±9118.34 | 1917.34±88.21 | 0.051±0.003 |
| Axodendritic | AB1 | 34 (7.31%) | 21 (4.52%) | 83910.48±13618.63 | 1273.60±117.31 | 0.046±0.004 |
| | AB2 | 56 (23.63%) | 21 (8.86%) | 69323.01±5321.07 | 1310.21±74.27 | 0.052±0.003 |
| | AB3 | 40 (9.88%) | 39 (9.63%) | 73100.66±9204.52 | 1302.21±92.32 | 0.042±0.003 |
| | AB4 | 35 (13.46%) | 37 (14.23%) | 81730.12±12154.13 | 1354.04±129.23 | 0.057±0.006 |
| | M17 | 23 (8.71%) | 41 (15.53%) | 191416.26±34376.21 | 2333.32±351.32 | 0.040±0.005 |

**Table S24.**
Ultrastructural data regarding the postsynaptic target in the *stratum radiatum* of CA1 per case. AS: asymmetric synapses; SAS: synaptic apposition surface; sem: standard error of the mean; SS: symmetric synapses.

The following significant differences were observed between cases:

- AB1-AB2 ($\chi^2$, p=5.799x$10^{-10}$)
  - Axospinous AS ($\chi^2$, p=7.427x$10^{-10}$)
  - Axodendritic AS ($\chi^2$, p=3.899x$10^{-8}$)
- AB1-AB4 ($\chi^2$, p=3.042x$10^{-7}$)
  - Axospinous AS ($\chi^2$, p=1.030x$10^{-6}$)
  - Axodendritic SS ($\chi^2$, p=1.099x$10^{-5}$)

- AB1-M17 ($\chi^2$, p=3.811x10$^{-6}$)
    - Axospinous AS ($\chi^2$, p=5.715x10$^{-5}$)
    - Axodendritic SS ($\chi^2$, p=6.787x10$^{-7}$)
- AB2-AB3 ($\chi^2$, p=2.195x10$^{-5}$)
    - Axospinous AS ($\chi^2$, p=0.0002)
    - Axodendritic AS ($\chi^2$, p=5.712x10$^{-6}$)
- AB2-M17 ($\chi^2$, p=3.466x10$^{-5}$)
    - Axodendritic AS ($\chi^2$, p=4.617x10$^{-6}$).

| *STRATUM LACUNOSUM-MOLECULARE* | | | | | | |
|---|---|---|---|---|---|---|
| | | No. AS | No. SS | SAS Area of AS ($nm^2$; mean±sem) | SAS Perimeter of AS (nm; mean±sem) | SAS Curvature of AS (mean±sem) |
| Axospinous | AB1 | 257 (76.95%) | 6 (1.80%) | 83065.94±3321.38 | 1404.69±40.24 | 0.045±0.002 |
| | AB2 | 69 (42.33%) | 0 (0.00%) | 108918.10±8387.64 | 1668.96±97.55 | 0.046±0.004 |
| | AB3 | 220 (59.30%) | 11 (2.96%) | 80266.24±3551.68 | 1437.99±41.37 | 0.047±0.003 |
| | AB4 | 132 (50.57%) | 9 (3.45%) | 109428.06±6696.00 | 1553.85±65.44 | 0.042±0.003 |
| | M17 | 82 (39.23%) | 5 (2.39%) | 108727.58±6012.23 | 1770.74±85.69 | 0.037±0.002 |
| Axodendritic | AB1 | 40 (11.98%) | 31 (9.28%) | 171377.95±19122.35 | 2163.14±182.35 | 0.049±0.006 |
| | AB2 | 54 (33.13%) | 40 (24.54%) | 167765.83±13316.03 | 2037.10±102.68 | 0.042±0.005 |
| | AB3 | 86 (23.18%) | 54 (14.56%) | 125214.64±10453.62 | 1786.71±99.75 | 0.050±0.004 |
| | AB4 | 69 (26.44%) | 51 (19.54%) | 183806.73±14303.31 | 2233.71±125.27 | 0.046±0.003 |
| | M17 | 51 (24.40%) | 71 (33.97%) | 149766.91±11574.26 | 1987.54±108.43 | 0.041±0.004 |

**Table S25.**
Ultrastructural data regarding the postsynaptic target in the *stratum radiatum* of CA1 per case. AS: asymmetric synapses; SAS: synaptic apposition surface; sem: standard error of the mean; SS: symmetric synapses.

The following significant differences were observed between cases:

- AB1-AB2 ($\chi^2$, p=1.000x$10^{-17}$)
    - Axospinous AS ($\chi^2$, p=7.100x$10^{-15}$)
    - Axodendritic AS ($\chi^2$, p=4.538x$10^{-8}$)
    - Axodendritic SS ($\chi^2$, p=1.393x$10^{-6}$)
- AB1-AB3 ($\chi^2$, p=1.221x$10^{-5}$)
    - Axospinous AS ($\chi^2$, p=5.428x$10^{-7}$)

    - Axodendritic AS ($\chi^2$, p=0.0001)
- AB1-AB4 ($\chi^2$, p=8.884x10$^{-10}$)
    - Axospinous AS ($\chi^2$, p=2.774x10$^{-11}$)
    - Axodendritic AS ($\chi^2$, p=9.745x10$^{-6}$)
    - Axodendritic SS ($\chi^2$, p=9.635x10$^{-5}$)
- AB1-M17 ($\chi^2$, p=1.000x10$^{-17}$)
    - Axospinous AS ($\chi^2$, p=1.000x10$^{-17}$)
    - Axodendritic AS ($\chi^2$, p=0.0002)
    - Axodendritic SS ($\chi^2$, p=2.737x10$^{-13}$)
- AB2-AB3 ($\chi^2$, p=8.108x10$^{-5}$)
    - Axospinous AS ($\chi^2$, p=0.0003)
- AB3-M17 ($\chi^2$, p=2.016x10$^{-7}$)
    - Axospinous AS ($\chi^2$, p=4.169x10$^{-6}$)
    - Axodendritic SS ($\chi^2$, p=1.011x10$^{-7}$).

| *STRATUM ORIENS* | | | | | | | | |
|---|---|---|---|---|---|---|---|---|
| | | No. AS | No. SS | % AS | % SS | SAS Area of AS ($nm^2$; mean±sem) | SAS Perimeter of AS (nm; mean±sem) | SAS Curvature of AS (mean±sem) |
| Macular | AB1 | 329 | 14 | 82.46% | 70.00% | 71579.24±1967.39 | 1193.81±18.81 | 0.043±0.001 |
| | AB2 | 864 | 40 | 89.91% | 83.33% | 59814.16±1365.19 | 1115.07±13.42 | 0.042±0.001 |
| | AB3 | 393 | 33 | 88.12% | 82.50% | 85542.47±3569.38 | 1410.09±38.35 | 0.044±0.001 |
| | AB4 | 252 | 13 | 72.83% | 86.67% | 84867.52±3346.80 | 1407.83±33.93 | 0.048±0.002 |
| | M17 | 442 | 33 | 89.11% | 76.74% | 69102.99±1881.60 | 1288.68±21.82 | 0.037±0.001 |
| Horseshoe-shaped | AB1 | 17 | 6 | 4.26% | 30.00% | 183219.77±16801.10 | 3085.08±246.51 | 0.082±0.014 |
| | AB2 | 29 | 5 | 3.02% | 10.42% | 182127.32±9673.00 | 2746.61±128.61 | 0.076±0.008 |
| | AB3 | 14 | 3 | 3.14% | 7.50% | 137568.89±23446.87 | 2325.37±398.38 | 0.049±0.006 |
| | AB4 | 9 | 1 | 2.60% | 6.67% | 167379.41±26519.90 | 2699.52±397.93 | 0.060±0.007 |
| | M17 | 17 | 7 | 3.43% | 16.28% | 185560.52±20627.61 | 3079.68±303.30 | 0.059±0.011 |
| Perforated | AB1 | 50 | 0 | 12.53% | 0.00% | 197825.31±10973.68 | 2395.02±119.56 | 0.054±0.003 |
| | AB2 | 62 | 1 | 6.45% | 2.08% | 208917.63±9628.77 | 2658.75±120.75 | 0.055±0.004 |
| | AB3 | 26 | 3 | 5.83% | 7.50% | 151181.17±26095.46 | 2212.36±363.18 | 0.054±0.009 |
| | AB4 | 80 | 1 | 23.12% | 6.67% | 163022.68±12781.49 | 2108.82±114.20 | 0.054±0.003 |
| | M17 | 32 | 3 | 6.45% | 6.98% | 223945.46±16074.55 | 2756.46±134.44 | 0.058±0.005 |
| Fragmented | AB1 | 3 | 0 | 0.75% | 0.00% | 229404.54±44724.53 | 2978.62±827.78 | 0.047±0.003 |
| | AB2 | 6 | 2 | 0.62% | 4.17% | 187914.30±42592.67 | 1978.91±285.29 | 0.131±0.027 |
| | AB3 | 13 | 1 | 2.91% | 2.50% | 221662.67±37955.74 | 2365.06±281.85 | 0.087±0.022 |
| | AB4 | 5 | 0 | 1.45% | 0.00% | 253460.37±50951.13 | 2519.40±548.65 | 0.127±0.049 |
| | M17 | 5 | 0 | 1.01% | 0.00% | 222988.12±54432.34 | 2871.76±773.04 | 0.104±0.027 |

**Table S26.**

Ultrastructural data regarding the shape of the synaptic junction in the *stratum oriens* of CA1 per case. AS: asymmetric synapses; SAS: synaptic apposition surface; sem: standard error of the mean; SS: symmetric synapses.

The following significant differences were observed between cases (all non-macular synaptic types were grouped into a single category for contingency analysis):

- AB1-AB2:
  - AS Macular-AS Non-macular ($\chi^2$, p=0.0003)
- AB2-AB4:
  - AS Macular-AS Non-macular ($\chi^2$, p=$3.300\text{x}10^{-15}$)
- AB3-AB4
  - AS Macular-AS Non-macular ($\chi^2$, p=$6.503\text{x}10^{-8}$)
- AB4-M17
  - AS Macular-AS Non-macular ($\chi^2$, p=$1.480\text{x}10^{-9}$).

| *STRATUM PYRAMIDALE* (DEEP) | | | | | | | | |
|---|---|---|---|---|---|---|---|---|
| | | No. AS | No. SS | % AS | % SS | SAS Area of AS ($nm^2$; mean±sem) | SAS Perimeter of AS (nm; mean±sem) | SAS Curvature of AS (mean±sem) |
| Macular | AB1 | 449 | 37 | 85.04% | 82.22% | 78730.15±2027.51 | 1283.49±19.06 | 0.042±0.001 |
| | AB2 | 1070 | 85 | 91.61% | 80.58% | 68976.39±1583.69 | 1209.09±15.29 | 0.041±0.001 |
| | AB3 | 598 | 20 | 86.17% | 51.28% | 73778.67±2017.49 | 1294.03±20.01 | 0.050±0.001 |
| | AB4 | 532 | 38 | 75.14% | 82.61% | 68926.88±1552.90 | 1257.46±16.92 | 0.050±0.001 |
| | M17 | 645 | 37 | 85.89% | 77.08% | 76324.38±1742.54 | 1316.01±17.47 | 0.039±0.001 |
| Horseshoe-shaped | AB1 | 31 | 3 | 5.87% | 6.67% | 149675.87±16501.34 | 2678.37±173.76 | 0.063±0.007 |
| | AB2 | 29 | 13 | 2.48% | 12.62% | 174669.50±13105.00 | 2826.08±170.45 | 0.075±0.009 |
| | AB3 | 22 | 10 | 3.17% | 25.64% | 215452.94±23446.74 | 3247.15±244.84 | 0.101±0.014 |
| | AB4 | 27 | 3 | 3.81% | 6.52% | 180541.81±14825.42 | 3214.16±173.74 | 0.101±0.013 |
| | M17 | 32 | 10 | 4.26% | 20.83% | 209129.34±19899.44 | 3265.64±218.18 | 0.092±0.012 |
| Perforated | AB1 | 44 | 4 | 8.33% | 8.89% | 176386.67±11306.29 | 2244.43±111.14 | 0.055±0.003 |
| | AB2 | 58 | 6 | 4.97% | 5.83% | 232495.35±12747.71 | 2767.16±110.76 | 0.060±0.004 |
| | AB3 | 52 | 8 | 7.49% | 20.51% | 223263.55±12861.35 | 2704.16±141.77 | 0.070±0.008 |
| | AB4 | 134 | 5 | 18.93% | 10.87% | 198734.15±7530.39 | 2546.44±94.12 | 0.065±0.004 |
| | M17 | 69 | 1 | 9.19% | 2.08% | 197780.76±12455.03 | 2541.75±135.32 | 0.058±0.004 |
| Fragmented | AB1 | 4 | 1 | 0.76% | 2.22% | 277288.60±50954.51 | 2101.88±361.68 | 0.099±0.043 |
| | AB2 | 11 | 1 | 0.94% | 0.97% | 211369.76±32166.87 | 1956.33±217.05 | 0.084±0.016 |
| | AB3 | 22 | 1 | 3.17% | 2.56% | 315335.94±22539.18 | 2640.33±149.18 | 0.130±0.014 |
| | AB4 | 15 | 0 | 2.12% | 0.00% | 262814.40±20115.67 | 2632.68±170.81 | 0.155±0.024 |
| | M17 | 5 | 0 | 0.67% | 0.00% | 307082.60±56381.11 | 2512.75±474.32 | 0.134±0.019 |

**Table S27.**
Ultrastructural data regarding the shape of the synaptic junction in the deep part of the *stratum pyramidale* of CA1 per case. AS: asymmetric synapses; SAS: synaptic apposition surface; sem: standard error of the mean; SS: symmetric synapses.

The following significant differences were observed between cases (all non-macular synaptic types were grouped into a single category for contingency analysis):

- AB1-AB2:
  - AS Macular-AS Non-macular ($\chi^2$, p=7.211x10$^{-5}$)
- AB1-AB4:
  - AS Macular-AS Non-macular ($\chi^2$, p=1.920x10$^{-5}$)
- AB2-AB3:
  - AS Macular-AS Non-macular ($\chi^2$, p=0.0003)
- AB2-AB4:
  - AS Macular-AS Non-macular ($\chi^2$, p=1.000x10$^{-17}$)
- AB2-M17:
  - AS Macular-AS Non-macular ($\chi^2$, p=0.0001)
- AB3-AB4:
  - AS Macular-AS Non-macular ($\chi^2$, p=1.756x10$^{-7}$)
- AB4-M17:
  - AS Macular-AS Non-macular ($\chi^2$, p=2.077x10$^{-7}$).

| *STRATUM PYRAMIDALE* (SUP) | | | | | | | | |
|---|---|---|---|---|---|---|---|---|
| | | No. AS | No. SS | % AS | % SS | SAS Area of AS ($nm^2$; mean±sem) | SAS Perimeter of AS (nm; mean±sem) | SAS Curvature of AS (mean±sem) |
| Macular | AB1 | 871 | 29 | 81.94% | 93.55% | 57496.39±1054.48 | 1153.86±11.86 | 0.047±0.001 |
| | AB2 | 1149 | 34 | 92.51% | 79.07% | 65443.73±1391.25 | 1192.36±13.57 | 0.051±0.001 |
| | AB3 | 1061 | 24 | 89.92% | 80.00% | 65334.44±1252.92 | 1247.35±13.50 | 0.047±0.001 |
| | AB4 | 542 | 38 | 77.76% | 77.55% | 84802.11±1985.86 | 1381.46±18.63 | 0.052±0.001 |
| | M17 | 894 | 39 | 89.31% | 90.70% | 81656.02±1662.13 | 1406.95±17.33 | 0.044±0.001 |
| Horseshoe-shaped | AB1 | 73 | 1 | 6.87% | 3.23% | 119267.30±7495.94 | 2255.35±99.02 | 0.074±0.005 |
| | AB2 | 37 | 4 | 2.98% | 9.30% | 254061.00±15689.07 | 3359.05±143.48 | 0.096±0.006 |
| | AB3 | 47 | 4 | 3.98% | 13.33% | 205616.90±10852.09 | 3172.30±147.89 | 0.101±0.009 |
| | AB4 | 25 | 4 | 3.59% | 8.16% | 245836.20±24434.06 | 3676.85±294.94 | 0.075±0.009 |
| | M17 | 30 | 3 | 3.00% | 6.98% | 241140.40±22881.63 | 3684.42±308.37 | 0.115±0.014 |
| Perforated | AB1 | 105 | 1 | 9.88% | 3.23% | 130096.60±5612.62 | 1995.67±62.80 | 0.081±0.005 |
| | AB2 | 47 | 1 | 3.78% | 2.33% | 235325.60±10943.00 | 2797.07±117.30 | 0.084±0.009 |
| | AB3 | 43 | 0 | 3.64% | 0.00% | 233736.20±13618.83 | 2738.23±126.19 | 0.084±0.008 |
| | AB4 | 112 | 3 | 16.07% | 6.12% | 233271.20±8764.45 | 2805.01±91.58 | 0.067±0.003 |
| | M17 | 64 | 1 | 6.39% | 2.33% | 292952.60±16089.11 | 3463.83±186.25 | 0.075±0.006 |
| Fragmented | AB1 | 14 | 0 | 1.32% | 0.00% | 212851.50±26291.48 | 2314.15±253.01 | 0.138±0.026 |
| | AB2 | 9 | 4 | 0.72% | 9.30% | 203875.90±33946.89 | 2238.76±256.80 | 0.072±0.011 |
| | AB3 | 29 | 2 | 2.46% | 6.67% | 244916.90±17971.50 | 2370.95±138.85 | 0.117±0.012 |
| | AB4 | 18 | 4 | 2.58% | 8.16% | 324793.90±21657.34 | 2725.81±189.21 | 0.157±0.013 |
| | M17 | 13 | 0 | 1.30% | 0.00% | 347186.20±25769.52 | 3167.86±262.15 | 0.171±0.036 |

**Table S28.**

Ultrastructural data regarding the shape of the synaptic junction in the superficial part of the *stratum pyramidale* of CA1 per case. AS: asymmetric synapses; SAS: synaptic apposition surface; sem: standard error of the mean; SS: symmetric synapses.

The following significant differences were observed between cases (all non-macular synaptic types were grouped into a single category for contingency analysis):

- AB1-AB2:
  - AS Macular-AS Non-macular ($\chi^2$, p=1.900x$10^{-15}$)
- AB1-AB3:
  - AS Macular-AS Non-macular ($\chi^2$, p=1.613x$10^{-8}$)
- AB1-M17:
  - AS Macular-AS Non-macular ($\chi^2$, p=1.843x$10^{-6}$)
- AB2-AB4:
  - AS Macular-AS Non-macular ($\chi^2$, p=1.000x$10^{-17}$)
- AB3-AB4:
  - AS Macular-AS Non-macular ($\chi^2$, p=1.767x$10^{-13}$)
- AB4-M17:
  - AS Macular-AS Non-macular ($\chi^2$, p=1.496x$10^{-11}$).

| *STRATUM RADIATUM* | | | | | | | | |
|---|---|---|---|---|---|---|---|---|
| | | No. AS | No. SS | % AS | % SS | SAS Area of AS ($nm^2$; mean±sem) | SAS Perimeter of AS (nm; mean±sem) | SAS Curvature of AS (mean±sem) |
| Macular | AB1 | 568 | 23 | 87.95% | 92.00% | 68441.72±1880.37 | 1252.81±19.74 | 0.050±0.001 |
| | AB2 | 809 | 21 | 90.12% | 87.50% | 57905.52±1741.87 | 1126.16±17.11 | 0.053±0.001 |
| | AB3 | 946 | 36 | 90.34% | 92.31% | 59199.20±1234.64 | 1159.89±13.48 | 0.045±0.001 |
| | AB4 | 436 | 34 | 76.55% | 64.29% | 72845.49±2015.31 | 1241.70±18.57 | 0.056±0.002 |
| | M17 | 579 | 27 | 84.87% | 80.95% | 70869.88±1911.78 | 1245.05±18.57 | 0.036±0.001 |
| Horseshoe-shaped | AB1 | 34 | 0 | 5.06% | 0.00% | 122773.40±12538.54 | 2018.18±144.47 | 0.062±0.010 |
| | AB2 | 39 | 1 | 4.34% | 4.17% | 198204.30±12709.33 | 3219.84±214.07 | 0.070±0.005 |
| | AB3 | 43 | 1 | 4.05% | 2.56% | 183521.60±12386.53 | 2706.35±126.51 | 0.069±0.006 |
| | AB4 | 20 | 5 | 4.07% | 11.90% | 173804.50±20244.18 | 3084.70±303.97 | 0.074±0.008 |
| | M17 | 14 | 14 | 3.92% | 33.33% | 227324.70±23151.54 | 3243.59±206.94 | 0.078±0.011 |
| Perforated | AB1 | 36 | 1 | 5.51% | 4.00% | 127297.40±12183.28 | 1828.05±112.44 | 0.066±0.009 |
| | AB2 | 40 | 0 | 4.34% | 0.00% | 212574.60±13737.09 | 2659.22±148.73 | 0.086±0.001 |
| | AB3 | 31 | 0 | 2.85% | 0.00% | 194467.80±11623.65 | 2808.49±193.32 | 0.083±0.013 |
| | AB4 | 99 | 1 | 16.29% | 2.38% | 197019.10±8629.29 | 2524.29±96.59 | 0.071±0.004 |
| | M17 | 68 | 1 | 9.66% | 2.38% | 277764.10±16430.86 | 3370.50±180.99 | 0.074±0.006 |
| Fragmented | AB1 | 9 | 1 | 1.49% | 4.00% | 203063.30±48386.31 | 2197.56±345.52 | 0.098±0.044 |
| | AB2 | 9 | 2 | 1.19% | 8.33% | 220118.50±24204.66 | 2163.58±163.91 | 0.105±0.018 |
| | AB3 | 28 | 2 | 2.76% | 5.13% | 213947.90±20066.83 | 2211.08±178.18 | 0.088±0.012 |
| | AB4 | 17 | 2 | 3.09% | 4.76% | 332424.10±26011.06 | 3113.61±261.65 | 0.162±0.022 |
| | M17 | 11 | 0 | 1.54% | 0.00% | 377822.90±28842.89 | 3730.26±284.32 | 0.139±0.018 |

**Table S29.**
Ultrastructural data attending to the shape of the synaptic junction in the *stratum radiatum* of CA1 per case. AS: asymmetric synapses; SAS: synaptic apposition surface; sem: standard error of the mean; SS: symmetric synapses.

The following significant differences were observed between cases (all non-macular synaptic types were grouped into a single category for contingency analysis):

- AB-AB4:
  - AS Macular-AS Non-macular ($\chi^2$, p=1.673x10$^{-7}$)
- AB2-AB4:
  - AS Macular-AS Non-macular ($\chi^2$, p=1.242x10$^{-13}$)
- AB3-AB4:
  - AS Macular-AS Non-macular ($\chi^2$, p=2.300x10-$^{15}$)
- AB4-M17:
  - AS Macular-AS Non-macular ($\chi^2$, p=7.151x10$^{-6}$).

| *STRATUM LACUNOSUM-MOLECULARE* | | | | | | | | |
|---|---|---|---|---|---|---|---|---|
| | | No. AS | No. SS | % AS | % SS | SAS Area of AS ($nm^2$; mean±sem) | SAS Perimeter of AS (nm; mean±sem) | SAS Curvature of AS (mean±sem) |
| Macular | AB1 | 372 | 41 | 81.76% | 95.35% | 68113.92±2188.68 | 1206.36±20.98 | 0.045±0.002 |
| | AB2 | 453 | 24 | 84.99% | 60.00% | 75667.98±2563.19 | 1273.63±21.51 | 0.040±0.001 |
| | AB3 | 600 | 52 | 85.47% | 74.29% | 63852.28±1708.22 | 1212.36±16.96 | 0.045±0.001 |
| | AB4 | 313 | 65 | 72.96% | 92.86% | 78330.93±2507.34 | 1254.88±24.03 | 0.038±0.002 |
| | M17 | 422 | 76 | 83.90% | 81.72% | 74589.08±2372.56 | 1289.81±23.98 | 0.036±0.001 |
| Horseshoe-shaped | AB1 | 45 | 1 | 9.89% | 2.33% | 162197.29±11223.32 | 2649.59±120.03 | 0.053±0.007 |
| | AB2 | 28 | 12 | 5.25% | 30.00% | 201575.63±13044.70 | 2964.05±157.15 | 0.054±0.005 |
| | AB3 | 43 | 9 | 6.13% | 12.86% | 129287.65±7785.24 | 2422.88±108.91 | 0.046±0.003 |
| | AB4 | 40 | 4 | 9.32% | 5.71% | 164325.03±11131.14 | 2636.94±128.61 | 0.062±0.006 |
| | M17 | 33 | 11 | 6.56% | 11.83% | 144353.95±8840.51 | 2691.56±144.27 | 0.055±0.006 |
| Perforated | AB1 | 35 | 1 | 7.69% | 2.33% | 198469.66±13753.14 | 2441.51±133.31 | 0.049±0.004 |
| | AB2 | 47 | 3 | 8.82% | 7.50% | 245905.08±13748.10 | 2805.81±111.97 | 0.057±0.006 |
| | AB3 | 42 | 5 | 5.98% | 7.14% | 164625.20±12911.39 | 2339.99±148.91 | 0.060±0.006 |
| | AB4 | 62 | 1 | 14.45% | 1.43% | 225485.79±12755.30 | 2612.39±115.85 | 0.059±0.005 |
| | M17 | 36 | 3 | 7.16% | 3.23% | 171401.99±10472.22 | 2515.02±143.45 | 0.060±0.007 |
| Fragmented | AB1 | 3 | 0 | 0.66% | 0.00% | 198066.31±42515.28 | 2050.85±367.29 | 0.114±0.060 |
| | AB2 | 5 | 1 | 0.94% | 2.50% | 224196.15±11217.44 | 2356.51±210.70 | 0.067±0.017 |
| | AB3 | 17 | 4 | 2.42% | 5.71% | 177905.27±18503.35 | 2100.79±151.01 | 0.067±0.008 |
| | AB4 | 14 | 0 | 3.26% | 0.00% | 218888.89±28799.14 | 2026.17±236.32 | 0.072±0.007 |
| | M17 | 12 | 3 | 2.39% | 3.23% | 195513.30±19527.34 | 2460.70±209.08 | 0.063±0.013 |

**Table S30.**

Ultrastructural data regarding the shape of the synaptic junction in the *stratum lacunosum-moleculare* of CA1 per case. AS: asymmetric synapses; SAS: synaptic apposition surface; sem: standard error of the mean; SS: symmetric synapses.

The following significant differences were observed between cases (all non-macular synaptic types were grouped into a single category for contingency analysis):

- AB1-AB2:
  - SS Macular-AS Non-macular ($\chi^2$, p=0.0001)
- AB1-AB4:
  - AS Macular-AS Non-macular ($\chi^2$, p=0.0002)
- AB2-AB4:
  - AS Macular-AS Non-macular ($\chi^2$, p=$5.796x10^{-6}$)
  - SS Macular-AS Non-macular ($\chi^2$, p=$6.553x10^{-5}$)
- AB3-AB4:
  - AS Macular-AS Non-macular ($\chi^2$, p=$3.761x10^{-7}$)
- AB4-M17:
  - AS Macular-AS Non-macular ($\chi^2$, p=$5.416x10^{-5}$).

| Case | Sex | Age (years) | Cause of death | Postmortem delay (h) | Neuropathological diagnosis |
|---|---|---|---|---|---|
| AB1 | Male | 45 | Lung cancer | <1 | No neurological alterations |
| AB2 | Female | 53 | Pulmonary shock | 4 | No neurological alterations |
| AB3 | Male | 53 | Bladder carcinoma | 3.5 | No neurological alterations |
| AB4 | Female | 65 | Bone sarcoma | 4.5 | No neurological alterations |
| M17 | Male | 36 | Bronchopneumonia | 2.5 | No neurological alterations |

**Table S31.**
Clinical information regarding the human cases. None of the five subjects had recorded neurological or psychiatric alterations.
